# Iron and Heme Coordinate Erythropoiesis through HRI-Mediated Regulation of Protein Translation and Gene Expression

**DOI:** 10.1101/408054

**Authors:** Shuping Zhang, Alejandra Macias-Garcia, Jacob C. Ulirsch, Jason Velazquez, Vincent L. Butty, Stuart S. Levine, Vijay G. Sankaran, Jane-Jane Chen

**Affiliations:** Institute for Medical Engineering and Science, Massachusetts Institute of Technology, Cambridge, MA 02139, USA; Division of Hematology/Oncology, Boston Children’s Hospital, Harvard Medical School, Boston, MA 02115, USA; Department of Pediatric Oncology, Dana-Farber Cancer Institute, Harvard Medical School, Boston, MA 02115, USA; Broad Institute of Massachusetts Institute of Technology and Harvard, Cambridge, MA 02142, USA; Program in Biological and Biomedical Sciences, Harvard University, Cambridge, MA 02138, USA; BioMicro Center, Massachusetts Institute of Technology, Cambridge, MA 02139, USA

## Abstract

Iron and heme play central roles in red blood cell production. However, the mechanisms by which iron and heme levels coordinate erythropoiesis remain incompletely understood. HRI is a heme-regulated kinase that controls translation by phosphorylating eIF2α. Here, we investigate the global impact of iron, heme and HRI on protein translation *in vivo* in murine primary erythroblasts using ribosome profiling. By defining the underlying changes in translation during iron and HRI deficiencies, we validate known regulators of this process, including *Atf4*, and identify novel pathways such as co-regulation of ribosomal protein mRNA translation. Surprisingly, we found that heme and HRI pathways, but not iron-regulated pathways, mediate the major protein translational and transcriptional responses to iron deficiency in erythroblasts *in vivo* and thereby identify previously unappreciated regulators of erythropoiesis. Our genome-wide study uncovers the major impact of the HRI-mediated integrated stress response for the adaptation to iron deficiency anemia.

## Introduction

Iron deficiency anemia is estimated to affect one-third of the global population (Lopez et al. 2016). In addition to being key components of hemoglobin, the primary oxygen transport molecule, cellular iron and heme levels impact globin synthesis and red blood cell production. Specifically, globin is transcriptionally regulated by BACH1 (BTB Domain and CNC homolog 1) (Igarashi and Sun 2006) and regulated at the level of protein translation by HRI (heme-regulated eIF2α kinase) (Chen 2007), both of which are heme-sensing proteins.

During heme deficiency induced by dietary iron deficiency (ID) in mice, HRI is activated and phosphorylates the α subunit of eukaryotic initiation factor eIF2 (eIF2α) to inhibit translation of α-and β-globin mRNAs, so as to prevent proteotoxicity resulting from heme-free globin chains (Han et al. 2001). Meanwhile, phosphorylated eIF2α (eIF2αP) selectively enhances the translation of activating transcription factor 4 (*Atf4*) mRNA (Suragani et al. 2012, Chen 2014). This coordinated translational repression of general protein synthesis with the specific translational enhancement of *Atf4* mRNA by eIF2αP is termed the integrated stress response (ISR) (Harding et al. 2003). ISR is a universal response to several types of cellular stress (Chen 2014, Pakos-Zebrucka et al. 2016) initiated by the family of eIF2α kinase. Besides HRI, mammalian cells have three additional eIF2α kinases, which are expressed in distinct tissues to combat specific physiological stress. PKR responds to viral infection (Kaufman 2000) while GCN2 senses nutrient starvations (Hinnebusch 1996). PERK is activated by endoplasmic reticulum (ER) stress (Ron and Harding 2000). All four eIF2α kinases respond to oxidative and environmental stresses.

In the erythroid lineage, HRI expression increases during differentiation with higher expression in the hemoglobinized erythroblasts (Liu et al. 2008). Starting at the basophilic erythroblast stage, HRI is the predominant eIF2α kinase, expressed two orders of magnitude higher than the other three eIF2α kinases (Kingsley et al. 2013), and is responsible for over 90% of eIF2α phosphorylation (Liu et al. 2008). HRI-ISR signaling is necessary for effective erythropoiesis during ID (Han et al. 2001) and acts by reducing oxidative stress and promoting erythroid differentiation (Suragani et al. 2012, Zhang et al. 2018). Furthermore, HRI-ISR represses *in vivo* mTORC1 signaling activated by the elevated erythropoietin (Epo) levels during ID specifically in the erythroid lineage (Zhang et al. 2018). Thus, HRI coordinates two key translation-regulation pathways, eIF2αP and mTORC1 during ID. However, the exact molecular mechanisms by which iron and heme regulate erythropoiesis are incompletely understood.

While transcriptional regulation during erythropoiesis has been studied extensively (Kerenyi and Orkin 2010, An et al. 2014), much less is known about translational control of this process (Mills et al. 2016, Khajuria et al. 2018). Ribosome profiling (Ribo-seq) has emerged as a powerful tool to interrogate translation genome-wide (Ingolia et al. 2009). Here, we performed Ribo-seq and mRNA-seq in primary basophilic erythroblasts to investigate how *in vivo* translation is regulated by iron, heme and HRI in an unbiased manner to gain a global understanding of the molecular mechanisms governing erythropoiesis. We hypothesized that by globally surveying the landscape of translational and concomitant transcriptional changes occurring in the context of HRI deficiency either in iron replete (+Fe) or deficiency (−Fe) conditions, we could gain important insights into the mechanisms by which iron and heme can coordinate the process of erythropoiesis. Our results demonstrate that ISR of HRI-mediated translational regulation and subsequent ATF4-mediated gene expression is the most highly activated and critical pathway in developing basophilic erythroblasts during iron-restricted erythropoiesis.

## Results

### Overview of Ribo-seq and mRNA-seq data

Beginning at the basophilic erythroblast stage, erythropoiesis is finely regulated by iron and heme levels (Chiabrando, Mercurio, and Tolosano 2014, Muckenthaler et al. 2017). HRI and heme biosynthesis are upregulated, hemoglobin is actively synthesized (Figure 1A) (Liu et al. 2008, Chen 2014), and the majority of terminal erythroid gene expression changes have begun (Kerenyi and Orkin 2010, Ulirsch et al. 2014). Thus, basophilic erythroblasts (thereafter referred as EBs throughout for simplicity) from *Wt*+Fe, *Wt*−Fe, *Hri*^−/−^+Fe, and *Hri*^−/−^−Fe fetal livers (FLs) were used as sources to generate Ribo-seq and mRNA-seq libraries for genome-wide analysis of gene expression changes (Figure 1A).

**Figure 1.**
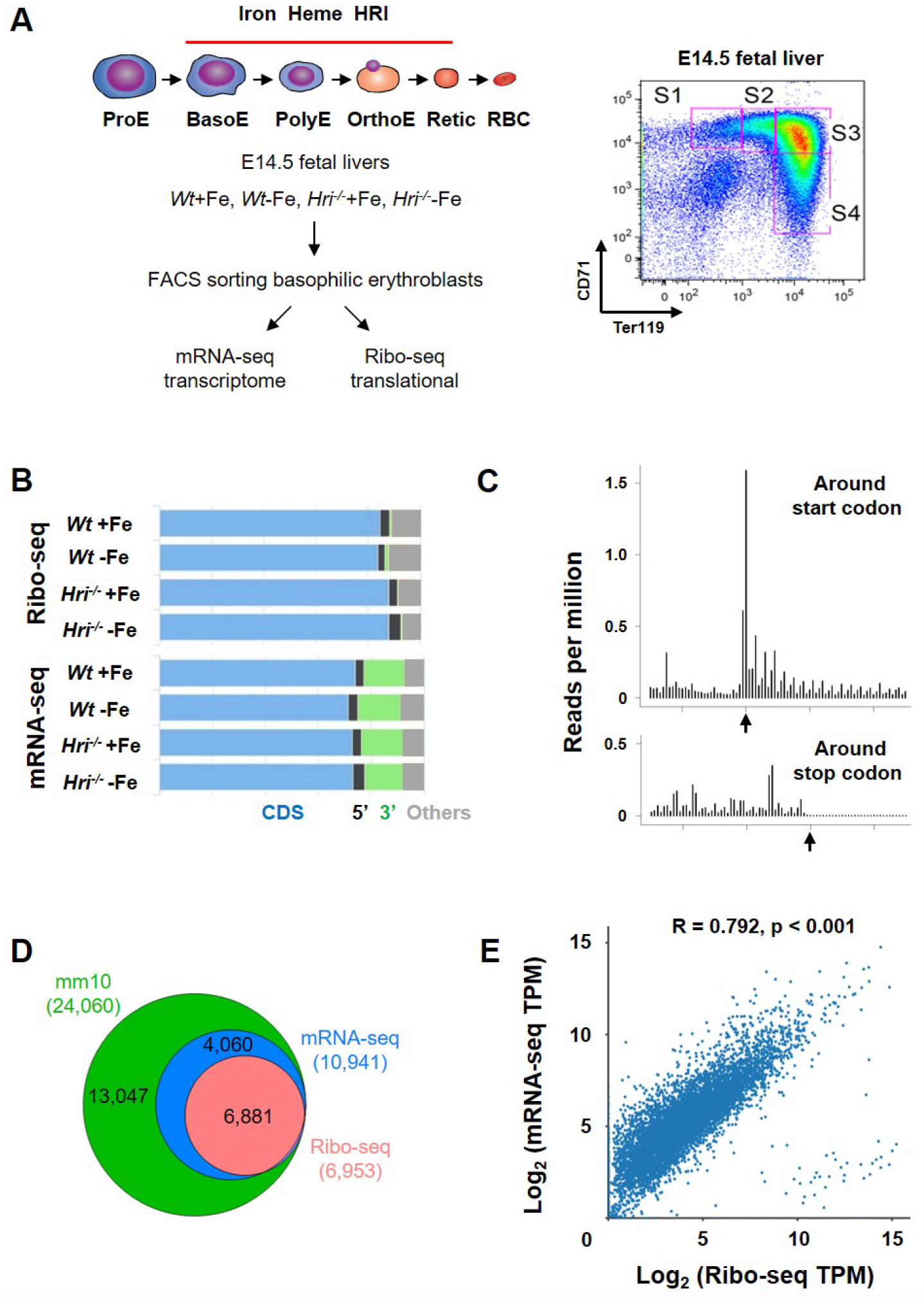
Overview of Ribo-seq and mRNA-seq data. **(A)** Illustration of experimental designs. Basophilic erythroblasts (EBs)(S3) from E14.5 FLs of *Wt*+Fe, *Wt*−Fe, *Hri*^−/−^+Fe and *Hri*^−/−^−Fe mice were sorted and subjected to the library preparations of Ribo-seq and mRNA-seq. **(B)** Distribution of the mapped reads from Ribo-seq and mRNA-seq from one replica. **(C)** A representative plot of triplet periodicity of Ribo-seq from *Wt*−Fe EBs. Arrow indicates the start and stop codons. **(D)** Gene coverages of Ribo-seq and mRNA-seq data in the mouse genome (UCSC, mm10). **(E)** Scatter plot and correlation analysis of log2-transformed TPM (transcript per million) of Ribo-seq and mRNA-seq data from *Wt*+Fe EBs. The following table supplement is available for Figure 1: Table supplement 1.

After applying standard protocols of quality control and preprocessing to remove rRNA and tRNA, we obtained 9.7-41.1 (median 26.2) million reads of Ribo-seq and 29.4-66.6 (median 42.5) million reads of mRNA-seq for subsequent mappings (Table supplement 1). As expected, most of the reads from both Ribo-seq (84-88%) and mRNA-seq (72-74%) were mapped to protein coding sequence (CDS) with some of reads mapped to the 5’ and 3’ UTR as well as other regions, mostly introns and the regions around transcription start sites (Figure 1B).

Our Ribo-seq data displayed excellent triplet periodicity, CDS enrichment, and limited 3’ UTR reads, validating the high quality of ribosome protected fragments (RPFs) that we obtained (Figure 1B-C). The proportions of reads mapping to 5’ UTRs of Ribo-seq data were similar to those from mRNA-seq (Figure 1B), in agreement with reports of pervasive translation outside of annotated CDSs (Ingolia et al. 2014). Overall, 62.9% of the expressed genes in mRNA-seq were detected in Ribo-seq, and therefore appeared to be actively translated in EBs (Figure 1D). As shown in Figure 1D, 99% of the mRNAs detected in Ribo-seq data (with reads greater than 25 in at least one of the conditions) were present in the mRNA-seq data, demonstrating the high quality of our Ribo-seq data and the correlation of RPFs and mRNAs (Figure 1E).

### Upregulation of *in vivo* translation of ISR mRNAs in *Wt* EBs compared to *Hri*^−/−^ EBs

317 mRNAs were significantly differentially translated between *Wt* and *Hri*^−/−^ EBs in +Fe condition (Figure 2A and Table supplement 2), supporting the role of HRI during normal fetal erythropoiesis. 25 differentially translated mRNAs were common under both +Fe and −Fe conditions and included the well-characterized ISR mRNAs, *Atf4, Ppp1r15a*, and *Ddit3* (Figure 2A-B and Figure supplement 1A). *Atf4* mRNA was the most differentially translated mRNA between *Wt* and *Hri*^−/−^ cells during ID (8.8-fold increase in translational efficiency (TE)), followed by *Ddit3* (8.1-fold) and *Ppp1r15a* (3.7-fold) mRNAs (Figure 2B and Table supplement 2). Each of these mRNAs contains upstream open reading frames (uORFs) in their 5’ UTR, and the use of which are upregulated by eIF2αP in cell lines under endoplasmic reticulum (ER) stress or amino acid starvation (Pavitt and Ron 2012). We observed HRI-dependent translation regulation of these mRNAs via ribosome occupancies in uORFs *in vivo* (Figure 2C-D and Figure supplement 1B-C). We also verified that changes in TE corresponded to change at *Atf4* protein levels in EBs (Figure 2E). In addition, we identified a potential novel candidate mRNA, *Brd2*, which may be regulated by uORFs via HRI-eIF2αP. TE of *Brd2* mRNA was higher in *Wt*−Fe EBs (2.34-fold) as compared to *Hri*^−/−^−Fe EBs (Figure supplement 2A). Ribosome occupancies were observed in putative uORFs at the 5’ UTR of *Brd2* mRNA (Figure supplement 2B). We validated the increased *Brd2* protein expression in *Wt*−Fe EBs compared to *Hri*^−/−^−Fe Ebs (Figure supplement 2C). Depletion of BRD2 expression was shown to inhibit terminal erythroid differentiation (Stonestrom et al. 2015).

**Figure 2.**
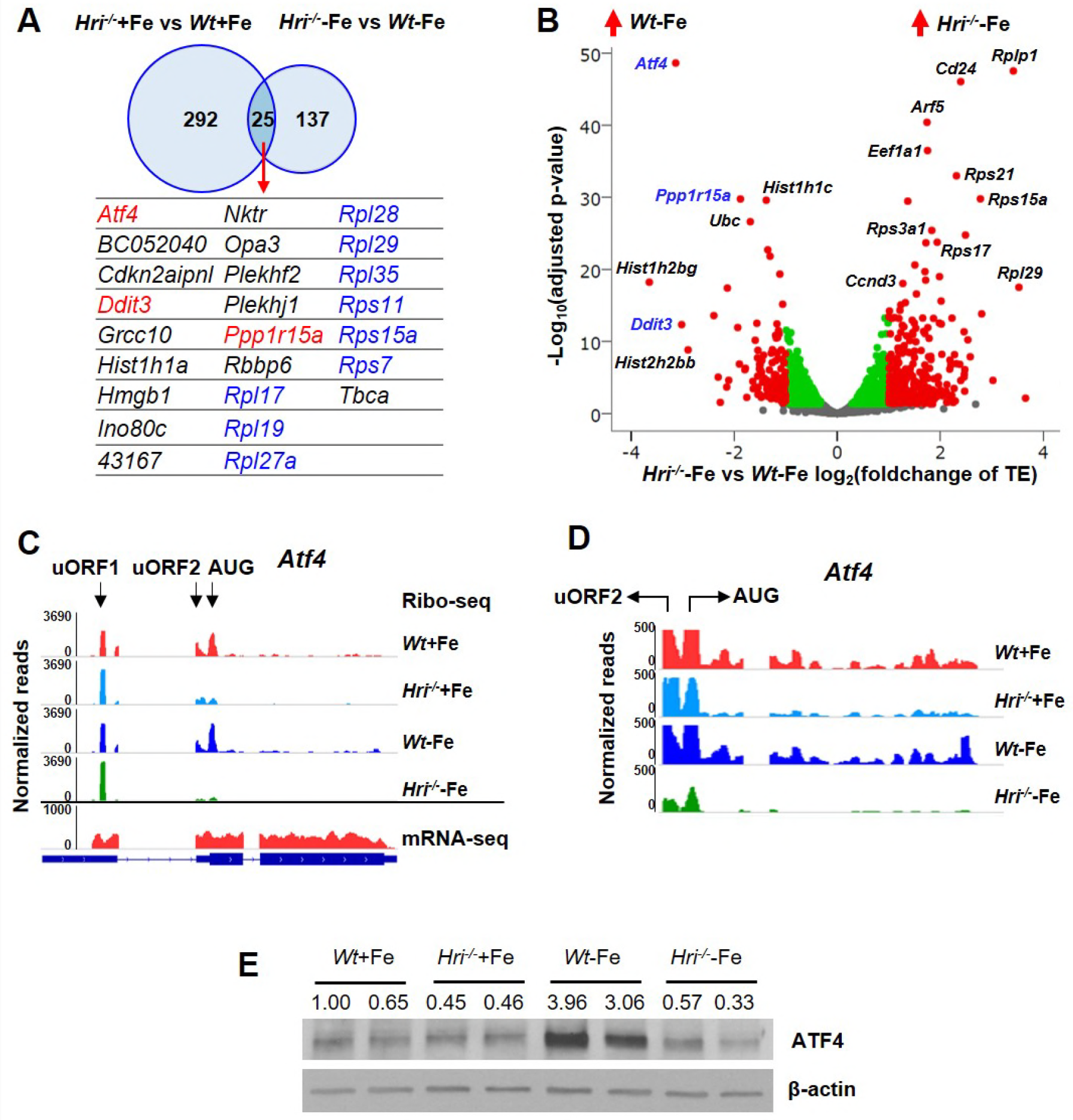
Differentially translated mRNAs in HRI and iron deficiencies. **(A)** The significantly differentially translated mRNAs between *Hri*^−/−^ and *Wt* EBs in +Fe or −Fe conditions. **(B)** The volcano plot of differentially translated mRNAs between *Hri*^−/−^−Fe and *Wt*−Fe EBs. Red dots on the positive end of X-axis indicate significantly differentially translated mRNAs upregulated in *Hri*^−/−^−Fe EBs while red dots on the negative end of X-axis indicate significantly differentially translated mRNAs upregulated in *Wt*−Fe EBs. Green and gray dots indicate not significantly differentially translated mRNAs. TE, translational efficiency. **(C)** Ribosome occupancies, as visualized using Integrative Genomics Viewer (IGV), of *Atf4* mRNA, with an enlarged view shown in **(D)**. **(E)** ATF4 protein expression in E14.5 FL cells. The following supplements are available for Figure 2: Figure supplement 1-2 and Table supplement 2-3.

Since *Hri* and *Atf4* are among the most highly expressed and efficiently translated mRNAs in *Wt* EBs (top 3%, Table 1), we investigated whether *Atf4* mRNA was poised for translational regulation by HRI. *Atf4* mRNA has two well-characterized uORFs in its 5’ UTR, including uORF1 which encodes for 3 amino acids and is translated regardless of stress and eIF2αP levels (Pavitt and Ron 2012). We observed that uORF1 had exceptionally high ribosome occupancy (5.7-fold and 21.3-fold higher than *eIF2s1* (eIF2α) and *Rps6* initiating AUGs, respectively, Table supplement 3). However, uORF2 and the canonical ORF of *Atf4* were poorly translated in *Hri*^−/−^ compared to *Wt* EBs under both +Fe and −Fe conditions (Figure 2C-D). Together, these data support the idea that *Atf4* mRNA is primed for translation by HRI in developing EBs.

**Table 1.** *Hri*, *Ppp1r15a* and *Atf4* mRNAs are highly expressed in basophilic erythroblasts.

| Gene Symbol | Rank | mRNA-seq |  |  |  | Ribo-seq |  |  |  |
| --- | --- | --- | --- | --- | --- | --- | --- | --- | --- |
|  |  | *Wt+Fe* | *Hri<sup>-/-</sup>+Fe* | *Wt+Fe* | *Hri<sup>-/-</sup>-Fe* | *Wt+Fe* | *Hri<sup>-/-</sup>+Fe* | *Wt+Fe* | *Hri<sup>-/-</sup>-Fe* |
| *Ftl1* | 6 | 10783.2 | 11396.0 | 7079.7 | 8726.3 | 324.5 | 416.6 | 297.1 | 558.0 |
| *Fth1* | 7 | 10619.8 | 10066.0 | 8485.8 | 10097.0 | 2308.3 | 2866.6 | 2447.9 | 4213.9 |
| *Alas2* | 8 | 9674.5 | 8269.8 | 7388.2 | 9954.1 | 5545.7 | 3962.9 | 4175.6 | 8487.4 |
| *Fech* | 18 | 4485.9 | 3937.3 | 3358.0 | 3991.4 | 1707.6 | 1647.4 | 1485.4 | 2958.5 |
| *Tfrc* | 57 | 2175.0 | 2158.2 | 2436.4 | 2456.6 | 1036.8 | 1436.1 | 2276.6 | 2483.1 |
| *Klf1* | 66 | 1964.8 | 1848.4 | 1706.2 | 1762.1 | 2687.8 | 2181.4 | 2335.0 | 2010.3 |
| *Nfe2* | 97 | 1380.9 | 1159.9 | 1077.5 | 1269.8 | 751.9 | 724.4 | 733.4 | 912.0 |
| *Eif2ak1 (Hri)* | 117 | 1223.4 | 1031.2 | 935.2 | 1053.3 | 566.5 | 461.4 | 631.6 | 488.1 |
| *Ppp1r15a* | 120 | 1222.1 | 1191.8 | 997.0 | 1451.1 | 751.4 | 254.8 | 278.6 | 158.1 |
| *Zfp1* | 121 | 1212.4 | 1143.1 | 1045.8 | 1039.6 | 89.6 | 53.3 | 128.1 | 100.5 |
| *Bcl2l1* | 156 | 989.2 | 745.0 | 619.7 | 865.8 | 225.7 | 137.1 | 96.4 | 130.9 |
| *Atf4* | 202 | 765.1 | 717.0 | 1124.9 | 729.9 | 648.2 | 176.8 | 466.7 | 57.7 |
| *Gata1* | 256 | 618.6 | 628.7 | 614.6 | 600.4 | 227.7 | 310.0 | 335.3 | 417.7 |
| *Foxo3* | 380 | 432.9 | 340.7 | 274.4 | 365.9 | 277.3 | 195.3 | 286.4 | 433.7 |
| *Ppp1r15b* | 908 | 176.3 | 184.4 | 177.9 | 165.2 | 15.0 | 20.7 | 35.7 | 23.2 |
| *Eif2s1* | 1345 | 117.0 | 132.1 | 132.2 | 106.0 | 52.3 | 86.5 | 87.1 | 80.8 |
| *Nfe2l2* | 1942 | 75.7 | 81.4 | 70.1 | 76.8 | 97.0 | 99.8 | 109.0 | 82.7 |
| *Ddit3* | 2016 | 72.0 | 65.5 | 165.0 | 83.7 | 112.8 | 10.3 | 111.5 | 13.1 |
| *Tfr2* | 2142 | 66.6 | 70.5 | 73.7 | 73.0 | 38.0 | 25.6 | 43.8 | 49.2 |
| *Bach1* | 2478 | 55.8 | 52.1 | 43.3 | 50.1 | 15.1 | 14.3 | 11.4 | 9.2 |
| *Grb10* | 3053 | 41.1 | 23.3 | 103.4 | 23.0 | 25.3 | 12.8 | 75.4 | 27.1 |
| *Ireb2 (Irp2)* | 3230 | 37.5 | 41.2 | 35.2 | 38.0 | 18.1 | 12.0 | 27.2 | 21.2 |
| *Atf5* | 4213 | 23.2 | 13.2 | 316.4 | 11.5 | 1.9 | 0.9 | 12.9 | 1.1 |
| *Eif2ak2* | 4520 | 19.7 | 20.2 | 14.3 | 21.2 | 16.3 | 11.7 | 7.4 | 13.7 |
| *Trib3* | 5603 | 10.0 | 1.7 | 127.2 | 4.1 | 0.5 | 0.3 | 20.3 | 0.8 |
| *Eif2ak3* | 6369 | 5.0 | 5.5 | 6.1 | 4.9 | 2.5 | 2.4 | 2.0 | 2.1 |
| *Aco1 (Irp1)* | 6449 | 4.5 | 5.6 | 3.8 | 5.7 | 1.3 | 1.7 | 3.2 | 4.7 |

\* mRNAs were ranked by the TPM of each mRNA from mRNA-seq data of *Wt+Fe* EBs. mRNAs of interests with respects to erythropoiesis, translation and ISR were compiled to highlight the abundance of *Hri*, *Ppp1r15a* and *Atf4* mRNAs.
Among the four eIF2α kinases, *Hri* (*Eif2ak1*) is most highly expressed in EBs, about 2 orders of the magnitude of *Pkr* (*Eif2ak2*) and *Perk* (*Eif2ak3*). *Eif2ak4* (*Gcn2*) was expressed at a level lower than the cut-off.
The following table supplement is available for Table 1: Table supplement 4.

In contrast, we observed limited changes in the translation of mRNAs containing iron-responsive elements (IREs) in 5’UTR or at the mRNA levels of TfR1 or DMT1 during ID or HRI deficiency (Table supplement 4 and Table 1). We found that mRNAs of *Alas2* and ferritin heavy chain (*Fth1*) were translated in high efficiency in both +Fe and −Fe conditions (Figure supplement 3 and Table 1). Unexpectedly, our genome-wide study further demonstrate that *Fth1* mRNA was translated preferentially (7-8 fold) over *Ftl1* mRNA (Figure supplement 3B and Table 1). Additionally, IRP1 and IRP2 mRNAs were expressed at low levels and poorly translated in EBs (Table 1).

These *in vivo* genome-wide results demonstrate that diet-induced ID, sufficient to induce anemia in mice, does not significantly affect gene expression through IRE/IRPs in primary developing EBs. Together, our results demonstrate that HRI is a master translational regulator of key ISR mRNAs *in vivo* in primary EBs especially during ID and suggest that HRI likely regulates the translation of these mRNAs via a uORF-mediated mechanism.

### Upregulation of translation of ribosomal protein mRNAs in *Hri*^−/−^−Fe EBs compared to *Wt*−Fe EBs

On the other hand, two translational pathways were upregulated in *Hri*^−/−^−Fe EBs as compared to *Wt*−Fe EBs (Figure 3A-B). First, we found that translation initiation complex formation was upregulated in *Hri*^−/−^−Fe cells (Figure 3B), consistent with the known function of HRI-eIF2αP in ternary and 43S complexes formation (Chen 2007). Second, a number of ribosomal protein mRNAs and other mTORC1 translational targets were upregulated in *Hri*^−/−^−Fe EBs compared to *Wt*−Fe EBs (Figure 2A-B and Figure 3B-C). Each of these ribosomal protein mRNAs have a 5’ terminal oligopyrimidine (TOP) motif in their 5’ UTR which permits regulation via mTORC1 signaling (Figure 3D) (Thoreen et al. 2012). Furthermore, we observed that *Eef1a1* and *Ccnd3*, which have been shown to be regulated by mTORC1 signaling (Thoreen et al. 2012), were more efficiently translated in *Hri*^−/−^−Fe cells compared to *Wt*−Fe EBs (Figure 2B). Overall, these genome-wide results indicate a role of the repressed eIF2αP and elevated mTORC1 signaling in *Hri*^−/−^−Fe EBs as compared to *Wt*−Fe EBs.

**Figure 3.**
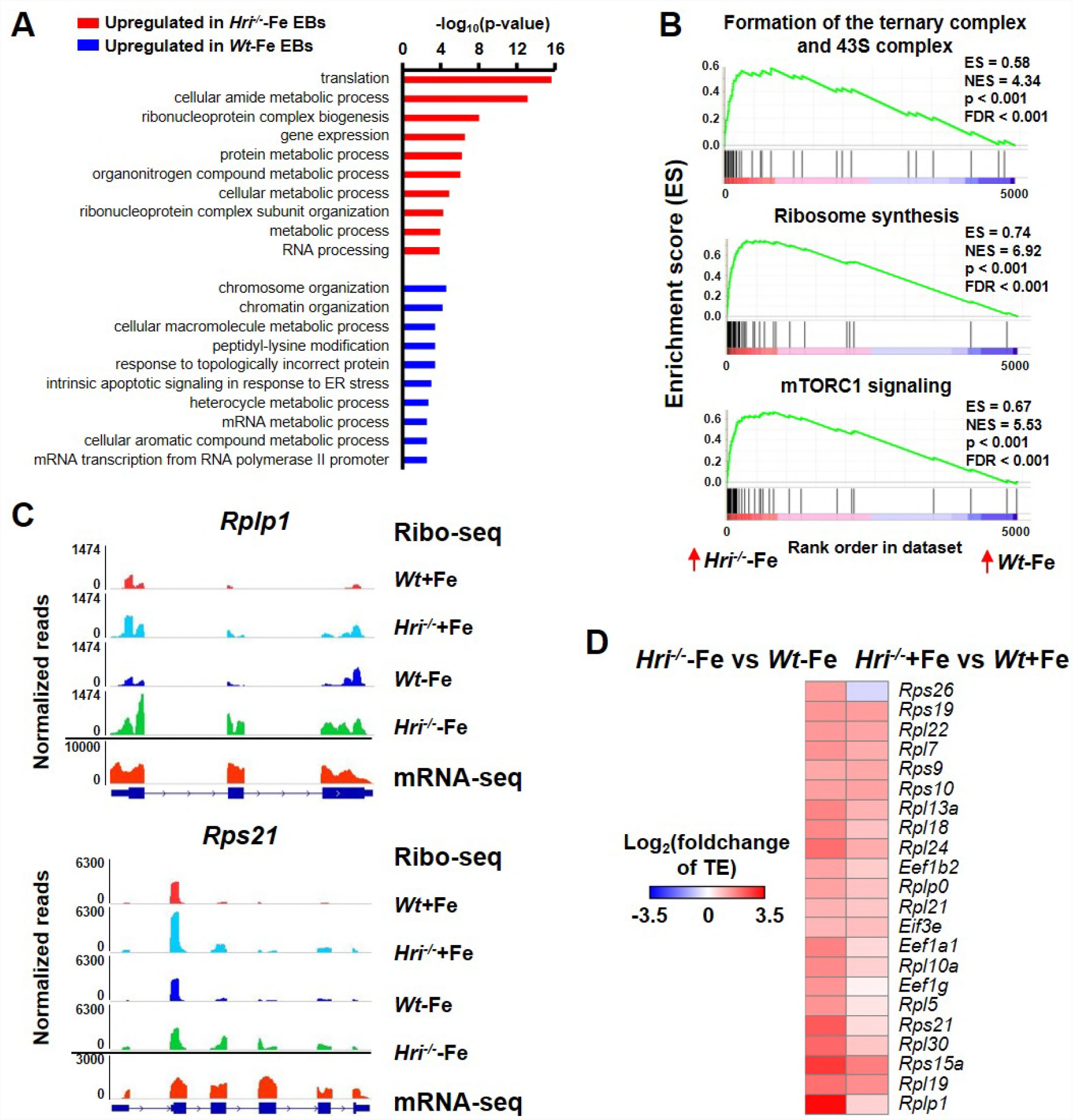
Analyses of the differentially translated mRNAs between *Wt* and *Hri*^−/−^ EBs. **(A)** Gene ontology analysis of the significantly differentially translated mRNAs between *Hri*^−/−^−Fe and *Wt*−Fe EBs. The top 10 most differentially changed biological processes in *Hri*^−/−^−Fe EBs as compared to *Wt*−Fe EBs are shown. **(B)** Three upregulated pathways in *Hri*^−/−^−Fe EBs as compared to *Wt*−Fe EBs revealed by Gene Set Enrichment Analysis. **(C)** IGV-illustration of ribosome occupancies of two representative ribosome mRNAs, *Rplp1* and *Rps21*. **(D)** The heatmaps of significantly differentially translated 5’TOP/TOP-like mRNAs, the mTORC1 translational targets. Positive values of Log2(foldchange of TE) indicate upregulated translation in *Hri-/-*−Fe EBs as compared to *Wt*−Fe EBs or in *Hri*^−/−^+Fe EBs as compared to *Wt*+Fe EBs. TE, translational efficiency.

### HRI-ATF4 mediated mRNA expression are most highly upregulated in ID

We next investigated the transcriptome impacts of ID and the role of HRI in mediating the cellular response to this stress. Analysis of mRNA-seq data revealed that there were substantially more genes that displayed significant differential expression between *Wt*−Fe and *Wt*+Fe EBs than between *Hri*^−/−^−Fe and *Hri*^−/−^+Fe EBs (232 vs 37, Figure 4A and Table supplement 5), demonstrating the near-absolute requirement for HRI in regulating the transcriptional response to ID. The majority of highly induced genes in *Wt*−Fe EBs compared to *Wt+*Fe EBs were ATF4 target genes, *Slc1a4, Atf5, Trib3, Asns, Shmt2, Pycr1 etc* (Figure 4B and 4D) (Pakos-Zebrucka et al. 2016). However, these genes were not up regulated in *Hri*^−/−^−Fe EBs (Figure 4C-D), indicating that HRI is required for activating ISR in ID (Figure 2). Several of these genes are involved in amino acid metabolism, which is the top biological process upregulated in *Wt*−Fe EBs compared to *Hri*^−/−^−Fe EBs as shown by gene ontology analysis (Figure supplement 4A). Furthermore, the expression levels of these genes were lower in *Hri*^−/−^ EBs than *Wt* EBs under +Fe condition (Figure supplement 4B), suggesting that HRI fine-tunes the ISR during iron replete erythropoiesis and thus has an important role even under normal conditions. Interestingly, some of these ATF4 target genes, most notably *Atf5, Trib3* and *Chac1*, are also Epo-stimulated genes (Figure 4E) (Singh et al. 2012), consistent with the interaction of HRI-ISR pathway and Epo signaling (Zhang et al. 2018).

**Figure 4.**
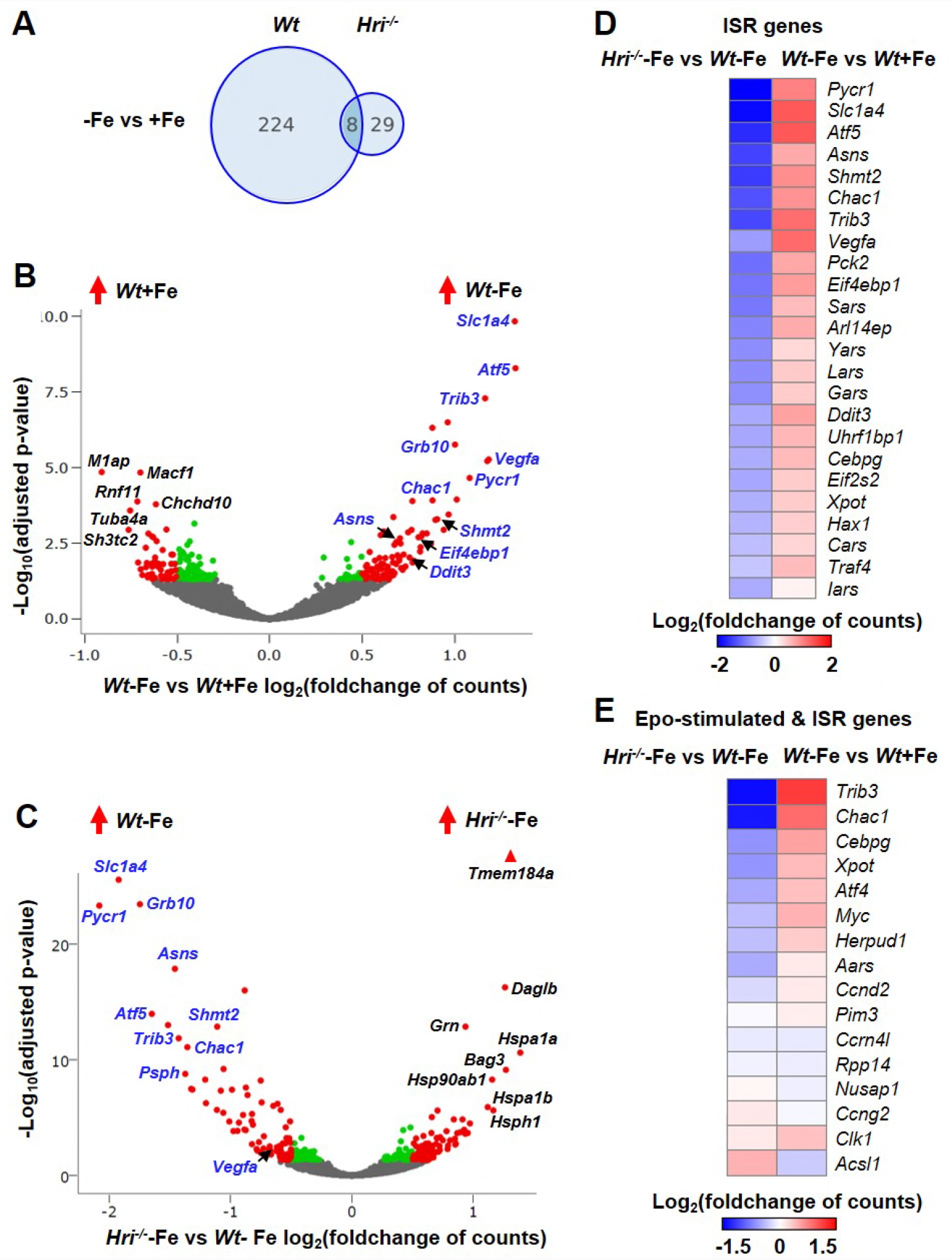
Differentially expressed mRNAs in HRI and iron deficiencies. **(A)** Numbers of the significantly differentially expressed mRNAs between −Fe and +Fe conditions of *Wt* or *Hri*^−/−^ EBs. Results were obtained from 3 biological replica. **(B)** Volcano plots of differentially expressed mRNAs between *Wt*−Fe and *Wt*+Fe or (**C)** between *Hri*^−/−^−Fe and *Wt*−Fe EBs. Red dots represent the significantly differentially expressed mRNAs. Green and gray dots indicate not significantly differentially expressed mRNAs. ATF4 target genes are labeled in blue. **(D)** Heatmaps of significantly differentially expressed ISR-target genes and **(E)** differentially expressed Epo-stimulated ISR-target genes between *Hri*^−/−^−Fe and *Wt*−Fe EBs or between *Wt*−Fe and *Wt*+Fe EBs. The following supplements are available for Figure 4: Figure supplement 4, Table supplement 5.

### Increased expression of chaperon mRNAs in *Hri*^−/−^−Fe EBs as compared to *Wt*−Fe EBs

The significantly highly upregulated genes in *Hri*^−/−^−Fe EBs, as compared to *Wt*−Fe EBs, were predominantly chaperone genes, *Hspa1a, Bag3, Hsp90ab1, Hspa1b* and *Hsph1* (Figure 4C), supporting the comprehensive upregulation of chaperone expression to mitigate globin inclusions in *Hri*^−/−^−Fe EBs. GO analysis also showed that processes and functions related to protein folding were enriched in *Hri*^−/−^−Fe EBs compared to *Wt*−Fe EBs (Figure supplement 4A). Moreover, *Hri*^−/−^−Fe EBs displayed gene expression characteristics of G2/M arrest as compared to *Wt*−Fe EBs (Figure supplement 4C), consistent with the increased cell percentages in G2/M of the cell cycle of splenic *Hri*^−/−^−Fe erythroid cells (Zhang et al. 2018).

### HRI-ISR necessary for fetal liver erythroid differentiation *ex vivo*

We have shown previously that HRI-ISR is activated in *ex vivo* FL differentiation system and that erythroid differentiation of *Hri*^−/−^ progenitors is impaired even under +Fe conditions (Suragani et al. 2012). Here, we show that there was no significant effect of HRI, eIF2αP, or ATF4 deficiencies on the growth and viability during expansion and up to 20 hours of erythroid differentiation (Figure 5A-B). However, *eIF2α Ala51/Ala51* (*AA*) erythroid precursors devoid of eIF2αP accumulated significant amounts of globin inclusions between 20-30 hours of the differentiation resulting in cell death with fragments of cell debris (Figure 5A). Similar observation was found in *Hri*^−/−^ erythroid precursors (Figure 5A), but less severe since eIF2αP is completely absence in *AA* erythroid precursors while *Hri*^−/−^ cells still have a low level of eIF2αP. These results underscore the first and foremost function of HRI-eIF2αP in inhibiting globin mRNA translation to mitigate proteotoxicity during FL erythropoiesis (Figure 5A).

**Figure 5.**
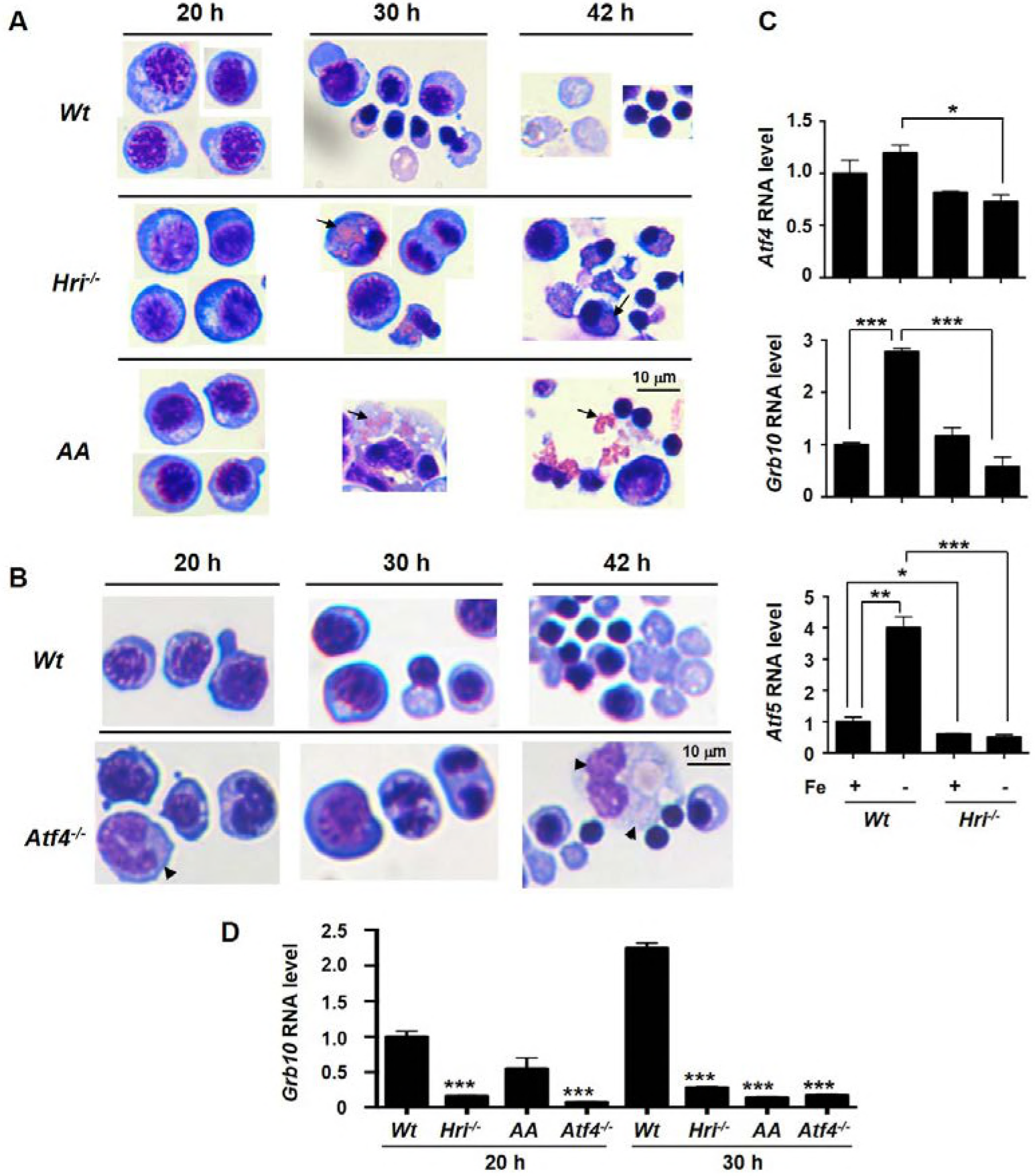
Impaired *ex vivo* FL differentiation and expression of *Atf4, Grb10, Atf5* RNAs in HRI-ISR defective erythroid cells. **(A-B)** *Ex vivo* erythroid differentiation from HRI, eIF2αP, and ATF4 deficient FL erythroid progenitors. The representative images of cytospin slides stained with May-Grunwald/Giemsa staining. Cells at 20, 30 and 42 hours of erythroid differentiation were shown. *AA*, universal *eIF2α Ala51/Ala51* knockin resulting in complete ablation of eIF2αP. Arrow indicates globin inclusions in (A) and arrowhead indicates myeloid cells in (B). Scale bar of 10 μm is indicated. Numbers of FL differentiation performed, n = 6 for *Wt* and *Hri*^−/−^; n = 4 for *AA* and n = 3 for *Atf4*^−/−^. **(C)** *Atf4, Grb10* and *Atf5* RNA expression in sorted basophilic EBs as illustrate in Figure 1A. Expression level in *Wt*+Fe EBs was defined as 1. (n = 3). ***p < 0.001. **(D)** *Grb10* RNA expression at 20 and 30 hours of *ex vivo* differentiation of *Wt, Hri*^−/−^, *AA* and *Atf4* FL erythroid progenitors. Expression level in *Wt* EBs at 20 h was defined as 1. For each time point, expression levels in *Hri*^−/−^, *AA* and *Atf4* EBs was compared to *Wt* EBs (n = 3). ***p < 0.001.

In contrast, *Atf4*^−/−^ FL erythroid cells did not suffer from proteotoxicity as would be expected due to the presence of functional HRI-eIF2αP in inhibiting globin mRNA translation (Figure 5B). However, differentiation of *Atf4*^−/−^ erythroid precursors at 30 and 42 hours was impaired (Figure 5B). In addition we also observed the presence of myeloid cells during erythroid differentiation of *Atf4*^−/−^ FL erythroid cells as compared to *Hri*^−/−^ or *AA* FL erythroid cells (Figure 5A-B). Together, these results demonstrate that both arms of ISR, inhibition of globin mRNA translation and enhanced translation of ATF4 mRNA are required for erythroid differentiation.

### *Grb10* required for late stage erythroid differentiation

Having mapped the genome-wide translational and transcriptional responses to ID, we set out to characterize the function of one of the top ATF4 target genes, growth factor receptor-bound protein 10 (*Grb10*) in erythropoiesis. Of all the genes that we identified as transcriptionally upregulated in *Wt*−Fe, but not *Hri*^−/−^−Fe EBs, *Grb10* (Figure 4B-C) was determined to be of particular interest since *Grb10* expression was not increased during ER stress (Harding et al. 2003), and it is therefore a novel gene that is specifically regulated by HRI-ISR under systemic ID. Furthermore, *Grb10* has been reported to be part of a feedback mechanism to inhibit growth factor-mediated mTORC1 signaling, such as insulin (Plasschaert and Bartolomei 2015) and stem cell factor (SCF) (Yan et al. 2016). We employed *ex vivo* FL differentiation to interrogate the function of *Grb10* in erythropoiesis.

First, we validated that *Grb10* and *Atf5*, but not *Atf4*, RNA expression was increased in *Wt* EBs, but not *Hri*^−/−^ EBs, during ID (Figure 5C). In addition, *Grb10* expression in *Wt* EBs was increased during *ex vivo* erythroid differentiation from 20 to 30 hours, and was greatly reduced in *Hri*^−/−^, *AA* and *Atf4*^−/−^ erythroblasts (Figure 5D). We prepared five shRNA recombinant retroviruses, all of which were able to knockdown *Grb10* RNA expression greater than 80% during the expansion phase. However, only one, shRNA_G3, was able to maintain persistent knockdown of *Grb10* RNA during differentiation (Figure 6A-B, Figure supplement 5 and Figure supplement 6A). Reduction of *Grb10* expression by shRNA_G3 both at RNA and protein levels (Figure 6B and 6F) increased cell numbers of differentiating erythroblasts (Figure 6C and Figure supplement 7A). This increase was first observed between 16 and 26 hours of differentiation (Figure 6C and Figure supplement 7A), in which Epo concentration was increased and SCF was withdrawn (Figure 6A). We also observed increased cyclin D3 expression (Figure 6F), indicating an increase of cells in the S phase of the cell cycle (Sankaran et al. 2012). Importantly, terminal erythroid differentiation was inhibited in *Grb10* knockdown cells as indicated by an accumulation of polychromatic erythroblasts, decrease of orthochromatic erythroblasts, and reduction of enucleation (Figure 6D-E, Figure supplement 6B-C and Figure supplement 7B-C). Thus, the increased proliferation but decreased terminal differentiation upon reduction of *Grb10* expression (Figure 6) recapitulates the hallmarks of ineffective erythropoiesis observed in *Hri*^−/−^ mice in ID (Han et al. 2001, Suragani et al. 2012, Zhang et al. 2018). Mechanistically, we observed that pAKTS473, a target of Epo signaling, was elevated in *Grb10* knockdown cells at 16 and 26 hours of differentiation (Figure 6F).

**Figure 6.**
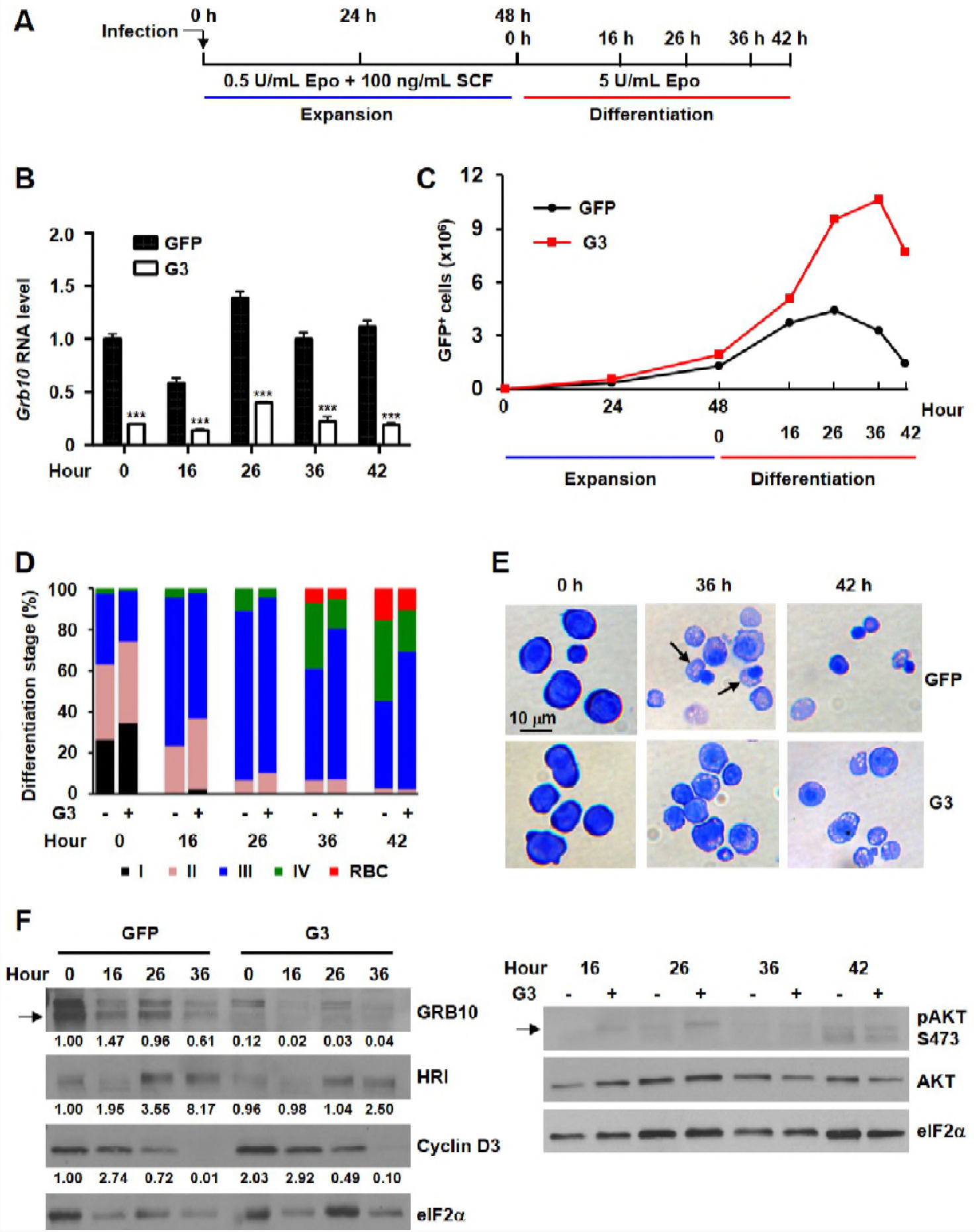
*Grb10* inhibits proliferation and promotes differentiation of erythroblasts.

**(A)** Designs of shRNA knockdown experiments using Lin^−^Ter119^−^CD71^−^ FL erythroid progenitors. Knockdown efficiency of *Grb10* RNA by shRNA_G3. *Grb10* expression in GFP control at 0 h is defined as 1. ***p < 0.001 (n = 3). **(C)** A representative proliferation of infected GFP^+^ cells. **(D)** Percentage of GFP^+^ cells at different erythroid differentiation stages and **(E)** Cell morphology of GFP control and *Grb10* knockdown cells. Scale bar of 10 μm is indicated. Arrow indicates orthochromatic erythroblast. **(F)** Epo and HRI signaling of GFP control and *Grb10* knockdown cells during differentiation. Three *Grb10* knockdown experiments were performed with similar results, and the results from a separate experiment are shown in the Figure supplement 7. GFP control, infected with retrovirus expressing GFP only; G3, infected with retroviruses expressing shRNA_G3. The following figure supplements are available for Figure 6: Figure supplement 5-7.

These results strengthen our global observation of a potential interaction between HRI-ISR and the Epo signaling pathway by showing that induction of *Grb10* by HRI-ISR serves as an important feedback mechanism for Epo signaling, resulting in reduced proliferation and thus promoting erythroid differentiation during stress erythropoiesis (Figure 7).

**Figure 7.**
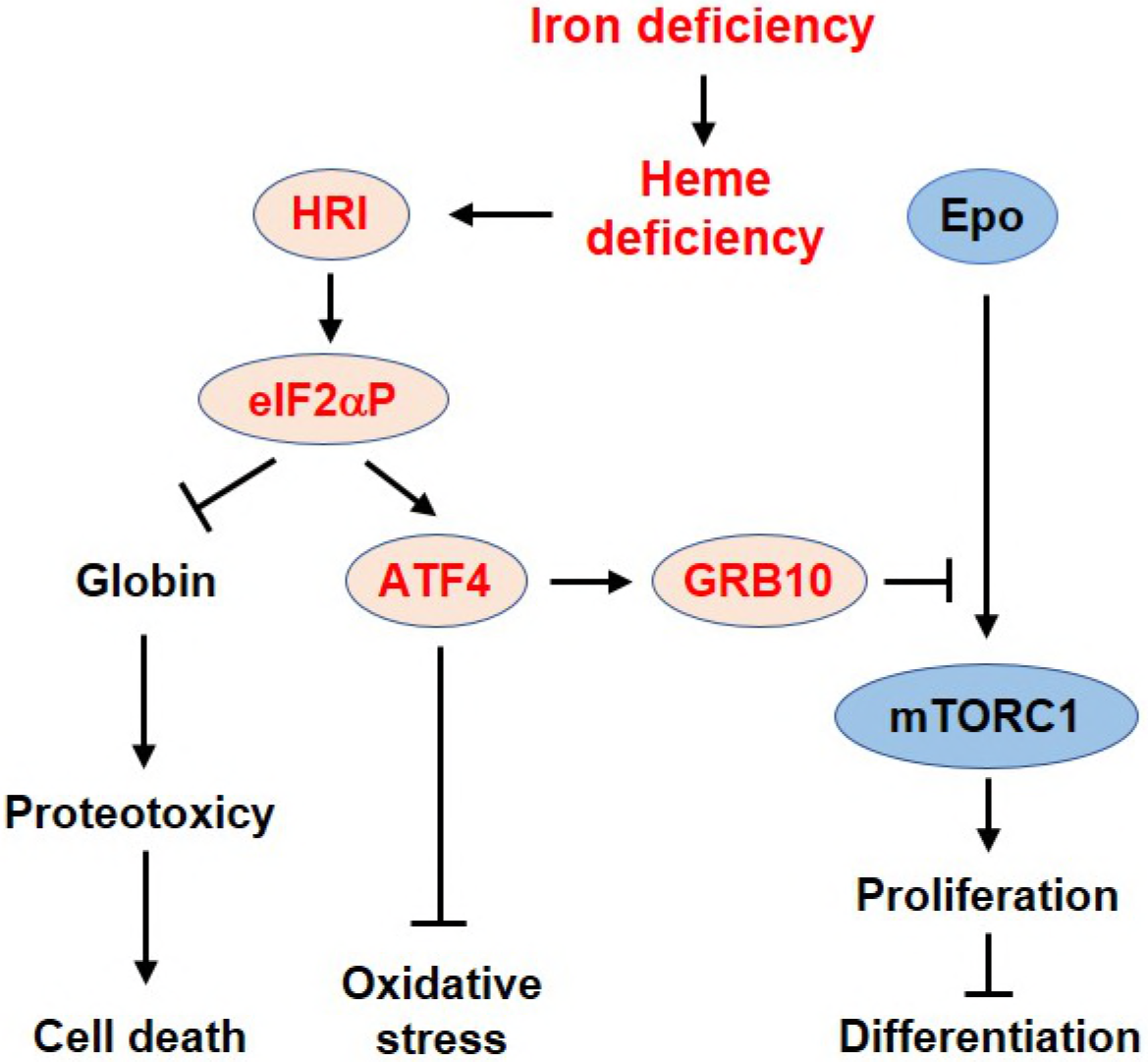
Proposed model of the regulation of erythropoiesis by heme and HRI-ISR pathway during ID. Iron is highly efficiently utilized for heme biosynthesis in developing EBs. There is no apparent IRE-mediated translational regulation in these cells during ID. Dietary induced systemic ID results in heme deficiency. HRI is activated in heme deficiency and phosphorylates its substrate eIF2α. The first and foremost function of eIF2αP is to inhibit globin mRNA translation to prevent proteotoxicity of unfolded heme-free globins. Secondly, eIF2αP also enhances the translation of *Atf4* mRNA via regulation by uORFs. *Atf4* mRNA is the most differentially translated mRNA between *Wt* and *Hri*^−/−^ EBs. Increased ATF4 expression then induces gene expression of an array of its target genes, including *Grb10*. Strikingly, expression of ATF4 target genes is most highly activated in ID and requires HRI. Thus, global genome-wide gene expression assessment of primary EBs *in vivo* reveals that HRI-ISR contributes most significantly in adaptation to ID. We have shown previously that HRI-ISR suppressed the mTORC1 signaling, which is activated by elevated Epo levels in ID (Zhang et al. 2018). Here, we provide evidence that GRB10 may be one of the molecules in repressing mTORC1 signaling. GRB10 is necessary to inhibit the proliferation and promote erythroid differentiation of erythroblasts upon stimulation by Epo.

## Discussion

While the specific role of HRI in translational regulation of globin and *Atf4* mRNAs in erythroid cells has been appreciated, an understanding of the global impact of HRI-mediated translation on erythropoiesis is lacking. Both iron and heme are necessary for terminal differentiation and can regulate protein translation, but the effectors that are critical for mediating responses to variable iron levels have remained undefined. Here, we report a global, unbiased assessment of iron, heme, and HRI-mediated translational and transcriptional alterations in primary EBs *in vivo*. Under our diet-induced ID in mice, which results in iron deficiency anemia, iron *per se* does not affect gene expression in EBs as translation of IRE-containing mRNAs are not altered during ID. Instead, our study demonstrates that heme is the major regulator of gene expression, at both the levels of translation and transcription, during erythropoiesis, as only limited changes of mRNA expression are observed in *Hri*^−/−^ EBs during ID. Thus, heme-regulated translation mediated by HRI is responsible for tuning gene expression responses to cellular heme levels during erythropoiesis in ID (Figure 7).

Our *in vivo* results in primary EBs are in agreement with the previous report of *Schranzhofer et al*, demonstrating that *Tfr1* mRNA stability, TfR1 protein expression, and bindings of IRP1 and IRP2 to IRE are not affected by iron depletion or iron supplement in differentiating erythroblasts from an immortalized cell line derived from FLs of E12.5 *p53*^−/−^ embryos (Schranzhofer et al. 2006). However, inhibition of heme synthesis restores iron-dependent ferritin protein synthesis. Therefore, iron is utilized very efficiently for heme synthesis such that developing erythroblasts is functionally iron deficient and IRE/IRP is sensing low iron state. *IRP2*^−/−^ mice develop microcytic anemia, demonstrate the essential role of IRP2 for erythropoiesis (Cooperman et al. 2005, Galy et al. 2005, Ghosh et al. 2008). While the neurodegenerative symptom of *IRP2*^−/−^ mice can be restored by IRP1 activation, this is not the case for anemia phenotype. Furthermore, IRP1 activity in erythroblasts was much lower than that from the brain, and was not activated by treatment with β-mercaptoethanol (Ghosh et al. 2008). Polycythemia developed in *IRP1*^−/−^ mice is not intrinsic to its deficiency in erythroid lineage but rather is indirectly resulting from the increased translation of renal HIF2α mRNA, which contains IRE in the 5’UTR, and subsequent increased Epo production (Ghosh et al. 2013, Anderson et al. 2013, Wilkinson and Pantopoulos 2013).

Recently, it has been shown that ferritin needs to be degraded in order to release its stored iron (Mancias et al. 2014). Nuclear receptor coactivator 4 (NCOA4) has been identified as a selective cargo receptor for lysosomal autophagic degradation of ferritin. Interestingly, NCOA4 binds only to FTH1, but not FTL1 (Mancias et al. 2015). Furthermore, NCOA4 is essential for erythroid differentiation (Mancias et al. 2015) and *NCOA4*^−/−^ mice developed severe microcytic hypochromic anemia when fed with iron-deficient diet due to the failure to release iron (Bellelli et al. 2016). This turnover of ferritin protein for the release of iron for heme and hemoglobin synthesis may explain the high rate of *Fth1* mRNA translation reported here to replenish ferritin protein homeostasis in EBs.

Most importantly, both *Hri* and *Atf4* mRNAs were abundantly expressed in EBs at the level on par with major erythroid transcriptional factors (*Klf1, Nfe2* and *Zfpm1*), and highly occupied by ribosomes, indicating that they are poised to respond to stimuli and stress during terminal erythroid differentiation. In contrast, *Bach1*, the other erythroid heme-sensing protein, was expressed at a level about 20-fold lower than *Hri* (Table 1). Our data further revealed that *Atf4*, the key transcription factor of the ISR pathway, and *Ppp1r15a* ISR mRNAs, are under exquisite HRI translational control in EBs, likely via mechanisms involving uORF translation.

*Ppp1r15a* gene encodes GADD34, which is a regulatory subunit of PPase1 bringing eIF2αP to PPase1 for dephosphorylation in order to reinitiate protein synthesis upon recovery from stress (Novoa et al. 2001, Kojima et al. 2003, Connor et al. 2001, Brush, Weiser, and Shenolikar 2003). Homeostasis of eIF2αP in Ter119^+^ erythroid cells is maintained by HRI and GADD34, both of which are necessary for erythropoiesis. Similar to *Hri*^−/−^ mice (Han et al. 2001), *Gadd34*^−/−^ mice, develop mild microcytic anemia with slight splenomegaly under normal conditions (Patterson et al. 2006). In contrast to *Hri*^−/−^ mice (Zhang et al. 2018), *GADD34*^−/−^ mice do not recover completely from iron deficiency anemia upon iron repletion due to persistent high levels of eIF2αP and inhibition of globin protein synthesis (Patterson et al. 2006). It is also noteworthy that *Atf4*^−/−^ mice develop transient anemia during fetal erythropoiesis (Masuoka and Townes 2002) and develop ineffective erythropoiesis in iron-deficient adult mice (Zhang et al. 2018). These prominent erythroid phenotypes of *Hri*^−/−^, *Atf4*^−/−^ and *Gadd34*^−/−^ mice are consistent with the results reported here on the highest global impact on the gene expression of these two genes by the activation of HRI-ISR signaling during iron-restricted erythropoiesis. It is to be noted that a recent study of ribosome profiling study was performed using immortalized FL cell line upon ER stress and not iron or HRI deficiency (Paolini et al. 2018), in contrast to our study of primary EBs directly from mice under iron replete and deficient conditions.

Our present study identifies several novel regulators of erythropoiesis, providing new insights into the regulation of stress erythropoiesis. We observed, at a global level, additional evidence of mTORC1 involvement in translational regulation in the absence of *Hri*, further supporting the role of HRI-ISR in repressing Epo-mTORC1 signaling to mitigate ineffective erythropoiesis during ID (Zhang et al. 2018). We provide supporting evidence that *Grb10* may be one of the ATF4 target genes in inhibiting Epo-mTORC1 signaling in ID. The increased proliferation of differentiating erythroblasts upon knockdown of *Grb10* reported here is consistent with the growth suppressor function of *Grb10*. *Grb10*, an imprinted gene, interacts with receptor tyrosine kinases and is a negative regulator of growth factor signaling (Plasschaert and Bartolomei 2015). Disruption of *Grb10* in mice results in the overgrowth and enhanced insulin signaling (Charalambous et al. 2003, Smith et al. 2007, Wang et al. 2007). In hematopoiesis, *Grb10* regulates the self-renewal and regeneration of hematopoietic stem cells (HSCs). *Grb10* deficient HSCs exhibit increased proliferation that is dependent on SCF-AKT/mTORC1 pathway (Yan et al. 2016), consistent with our *Grb10* knockdown results.

The increased proliferation of differentiating erythroblasts and impairment of terminal erythroid differentiation upon reduction of *Grb10* expression are the hallmarks of ineffective erythropoiesis, which characterizes diseases such as thalassemia and the myelodysplastic syndromes. We have shown earlier that HRI is activated in β-thalassemic erythroid cells and is necessary to reduce the severity of the disease symptoms in the mouse model (Han, Fleming, and Chen 2005). Thus, the HRI-ISR-GRB10 pathway may therefore be exploited for developing novel treatment for these diseases.

Induction of fetal hemoglobin (HbF) has been documented to ameliorate the symptoms of β-thalassemia and sickle cell anemia (Sankaran and Orkin 2013). HRI-ISR is necessary to reduce the severity of β-thalassemia syndrome in mouse model (Han, Fleming, and Chen 2005, Suragani et al. 2012). HRI-ISR has been shown to increase translation of fetal γ-globin mRNA by Hahn and Lowrey in human CD34+ cell culture through an unknown mechanism (Hahn and Lowrey 2013). Most recently, HRI was suggested to act as a repressor for HbF production by a domain-focused CRISPRA genome editing screen (Grevet et al. 2018). HRI depletion diminishes the expression of BCl11A mRNA, a repressor of γ-globin transcription, although the underlying mechanisms remain unknown. Interestingly, this HbF repressor action of HRI appears to be unique to human, and not observed in murine erythroid cells (Grevet et al. 2018). HRI deficiency did not result in significant changes of TE or levels of Bcl11A mRNA in our study report here (Table supplement 2 and Table supplement 5). The exact role of HRI in HbF production remains to be clarified.

In summary, our genome-wide study reveals the prominent contribution of HRI-ISR signaling in erythropoiesis and provides molecular insights on HRI-ISR-GRB10 signaling in the blunted Epo response during ID.

## Materials and Methods

### Animals and diet-induced iron deficiency

Mice were maintained at the Massachusetts Institute of Technology (MIT) animal facility, and all experiments were carried out using protocols approved by the Division of Comparative Medicine, MIT. *Hri*^−/−^, *Atf4*^−/−^ and universal *eIF2α Ala51/+* heterozygote knockin mice were as described previously (Han et al. 2001, Masuoka and Townes 2002, Scheuner et al. 2001). Diet-induced iron deficiency in mice was as previously described (Han et al. 2001, Zhang et al. 2018). Mice (8-12 weeks old) were mated for E13.5 or E14.5 fetal livers as sources of erythroid precursors. Under these iron deficient conditions, embryos were pale and anemic with decreased hematocrits in embryonic blood (Liu et al. 2008).

### Isolation of EBs, library preparations, DNA sequencing and genome-wide data analysis

EBs were sorted using anti-Ter119 and anti-CD71 antibodies by flow cytometry using FACS Aria (BD Biosciences, San Jose, CA) from E14.5 FLs of *Wt* and *Hri*^−/−^ mice maintained under +Fe or −Fe conditions (Figure 1A). In order to have sufficient EBs for Ribo-seq library, FLs from embryos of the same mother were pooled then sorted for EBs as one biological replica. Two (5 million cells each) and three biological replicas (1 million cells each) of each condition from separate mothers were collected for preparations of Ribo-seq and mRNA-seq libraries, respectively. The third replica of Ribo-seq using 3 million cells was unsuccessful likely due to lower cell numbers. All procedures of labeling and washing of cells for sorting were carried out at 4°C. Cells were sorted into tubes with 20% fetal bovine serum (FBS, Atlanta Biologicals, Norcross, GA) to help preserving cell integrity.

Ribo-seq libraries were prepared using ARTseq-Ribosome Profiling Kit (Illumina, San Diego, CA) as previously described (Guo et al. 2010, Ingolia, Lareau, and Weissman 2011) (see details in Method supplements). Total RNAs were extracted by RNeasy Plus kit (Qiagen, Germantown, MD) and polyA^+^ mRNAs were isolated using an Oligotex mRNA kit (Qiagen, Germantown, MD). mRNA-seq cDNA libraries were prepared by the MIT BioMicro Center. cDNA libraries of RPFs and mRNA were sequenced on HiSeq 2000 platform (Illumina, San Diego, CA) at the MIT BioMicro Center. After standard preprocessing and quality control analysis, reads were mapped to mouse genome mm10 (UCSC) followed by the downstream analyses (see details in Method supplements).

All Ribo-seq data and mRNA-seq described in this paper are available at the Gene Expression Ominibus (http://www.ncbi.nlm.nih.gov/geo/) under accession GSE119365.

### Enrichment of FL erythroid progenitors for *ex vivo* culture and differentiation

Erythroid progenitors from E13.5 FLs of *Wt*+Fe embryos were enriched by magnetic sorting using EasySep Magnet (StemCell Technologies, Vancouver, Canada) as described (Thom et al. 2014) (see details in Method supplements) and was illustrated in Figure supplement 5. Purified Lin^−^Ter119^−^CD71^−^ erythroid progenitors were cultured in expansion media as described (Thom et al. 2014) for two days at an initial cell density of 0.5 × 10^6^ cells per mL. Then, cells were washed and cultured in differentiation media at an initial cell density of 1 × 10^6^ cells per mL. Cells were collected for analysis at different time points as indicated in the Figure Legends and Figure 6A. Details of expansion media and differentiation media were described in Method supplements.

### Knockdown of *Grb10* expression in FL EBs by retroviral shRNAs

The knockdown of *Grb10* expression was performed by using recombinant retroviruses containing shRNA-expressing murine stem cell retroviral vector, MSCV-pgkGFP-U3-U6P-Bbs, a kind gift from the laboratory of Dr. Harvey F. Lodish (MIT). DNA sequences of five shRNA oligonucleotides (Table supplement 6) targeting different regions of *Grb10* mRNA were obtained from the Genetic Perturbation Platform of Broad Institute and synthesized by Integrated DNA Technologies. Preparations of the plasmid constructs and recombinant retroviruses were performed as described (Hu, Yuan, and Lodish 2014). Lin^−^Ter119^−^CD71^−^ erythroid progenitors enriched from *Wt*+Fe E13.5 FLs were infected with retroviruses as described (Thom et al. 2014). Cells were expanded for 48 hours after retroviral infections followed by differentiation up to 42 hours as indicated.

### Analysis of erythroid differentiation and cell proliferation

Erythroid differentiation was performed by flow cytometry using anti-Ter119 and anti-CD71 antibodies (BioLegend, San Diego, CA) as described (Zhang et al. 2018) as well as Draq5 florescent dye (BioLegend, San Diego, CA) for enucleation analysis on FACS LSR II (BD Biosciences, San Jose, CA). 4’,6-diamidino-2-phenylindole (DAPI, Roche Diagnostics, Basel, Switzerland) was used to exclude the dead cells. Data were analyzed with FlowJo (Tree Star, Ashland, OR). Erythroid differentiation was also analyzed by cell morphology on cytospin slides stained with May-Grunwald/Giemsa staining (Sigma-Aldrich, St. Louis, MO). Cell proliferation was determination by counting daily nucleated cells using crystal violet stain.

### RT-qPCR and Western blot analyses

Gene expression was performed by RT-qPCR and Western blot analyses as previously described (Zhang et al. 2018). Primers were listed in Table supplement 7. *Gapdh* was used as internal control for RT-qPCR. Antibodies used in Western blot were described in Table supplement 8. β-actin or eIF2α was used as a loading control for western blot.

### Statistical Analysis

Independent *t* test (two-tailed) was used to analyze the experimental data. Pearson correlation analysis was performed to determine the correlation coefficient. Data were presented in mean ± SE. *p < 0.05 was considered statistically significant. **p < 0.01; ***p < 0.001.

## Acknowledgments

This paper is dedicated to the memory of Irving M. London and his generous and inspiring mentorship. This work was supported by National Institute of Health Grant RO1 DK087984 (to J-JC), R01 DK103794 and R33 HL120791 (to V.G.S.).

## Authorship Contributions

S. Z., A. M-G., J. V. and J-J. C performed experiments. S. Z., J. C. U., V. G. S., V. L. B. and S. S. L. analyzed sequencing data. S. Z. and J-J. C. wrote the paper. A. M-G., J. C. U. and V. G. S. edited the paper.

## Disclosure of Conflicts of Interest

All authors declare no competing interests.

## Figure supplements

**Figure supplement 1.**
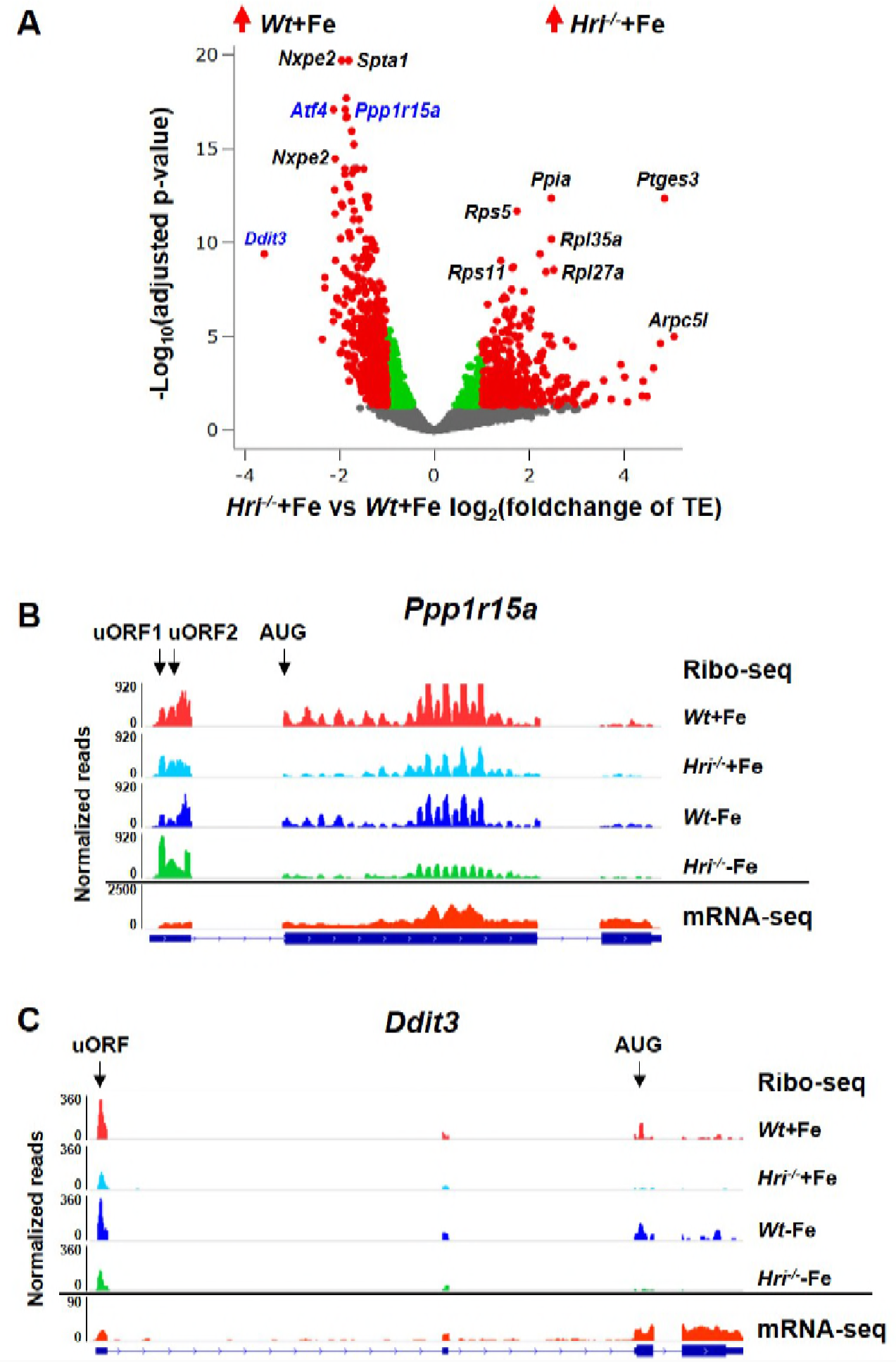
Translational regulation of ISR mRNAs by HRI. **(A)** Increased translation of ISR mRNAs in *Wt*+Fe EBs compared to *Hri*^−/−^+Fe EBs. Red dots on the positive end of X-axis indicate significantly differentially translated mRNAs which are upregulated in *Hri*^−/−^+Fe EBs as compared to *Wt*+Fe EBs while red dots on the negative end of X-axis indicate those upregulated in *Wt*+Fe EBs as compared to *Hri*^−/−^+Fe EBs. Green and gray dots indicate not significantly differentially translated mRNAs. Genes labeled in blue indicate ISR mRNAs. TE, translational efficiency. **(B-C)** IGV-illustration of ribosome occupancies of the *Ppp1r15a* and *Ddit3* mRNAs. Mapped reads of mRNA-seq data from *Wt*+Fe EBs were shown only since no significant change at mRNA levels was observed among four samples.

**Figure supplement 2.**
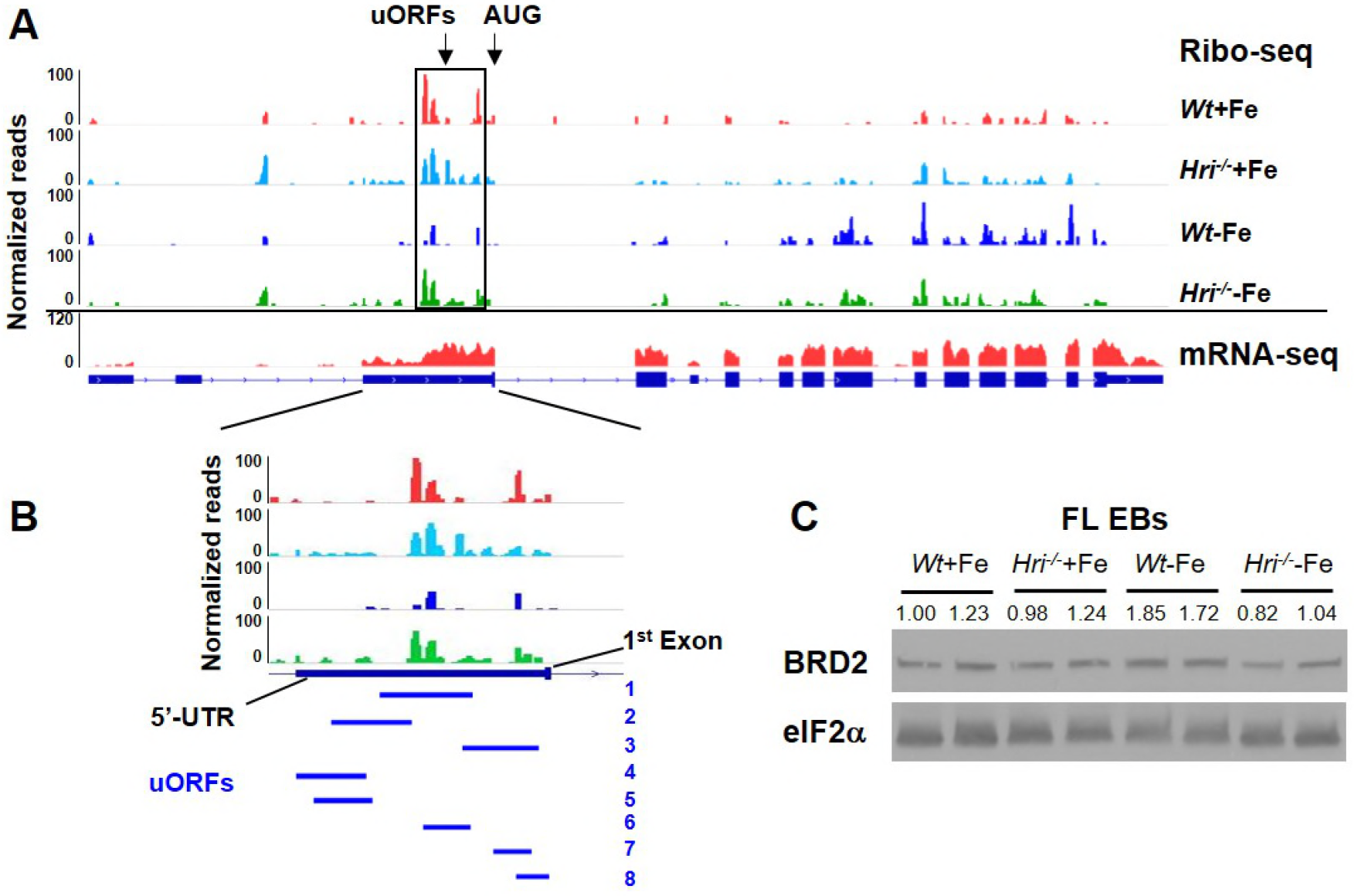
Putative translational regulation of *Brd2* mRNA by HRI in ID. **(A)** IGV-illustration of ribosome occupancies of *Brd2* mRNA. **(B)** The enlarged view of the region containing the putative uORFs which were indicated by the blue lines. Only the second half of the 5’ UTR sequence (−1269 to +26) was analyzed for the prediction of the presence of putative uORFs due to the long length of whole UTR and the low ribosome density on the first half of 5’ UTR. Eight uORFs were predicted, and ribosome occupancies were observed on all putative uORFs. **(C)** The protein levels of BRD2 in E14.5 FLs.

**Figure supplement 3.**
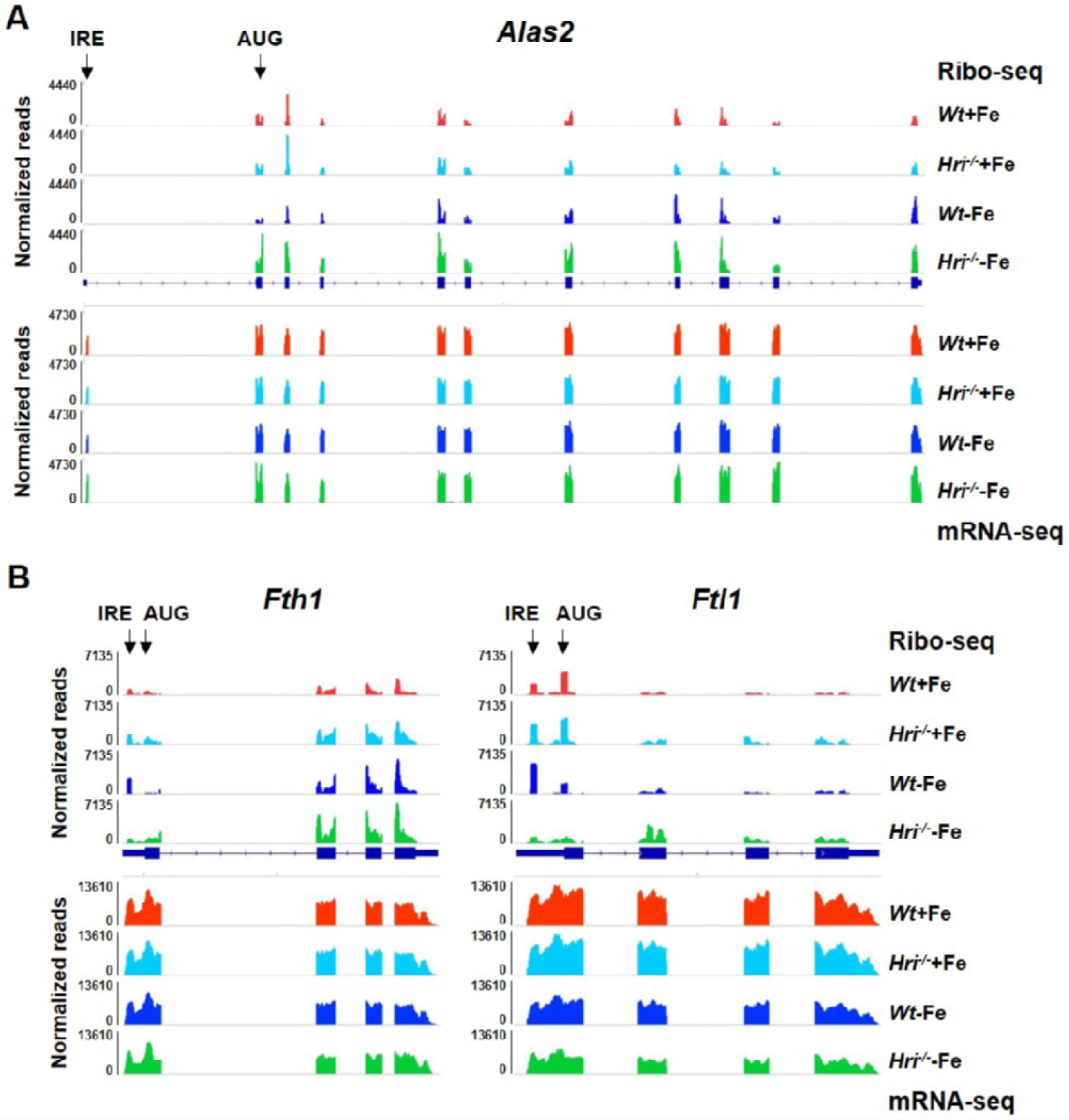
No significant alteration of translation of ALAS2 and ferritin mRNAs in ID. IGV-illustration of ribosome occupancies of **(A)** *Alas2*, and **(B)** *Fth1* and *Ftl1* mRNAs. These mRNAs contain IRE in 5’UTR and can be regulated translationally by IRPs.

**Figure supplement 4.**
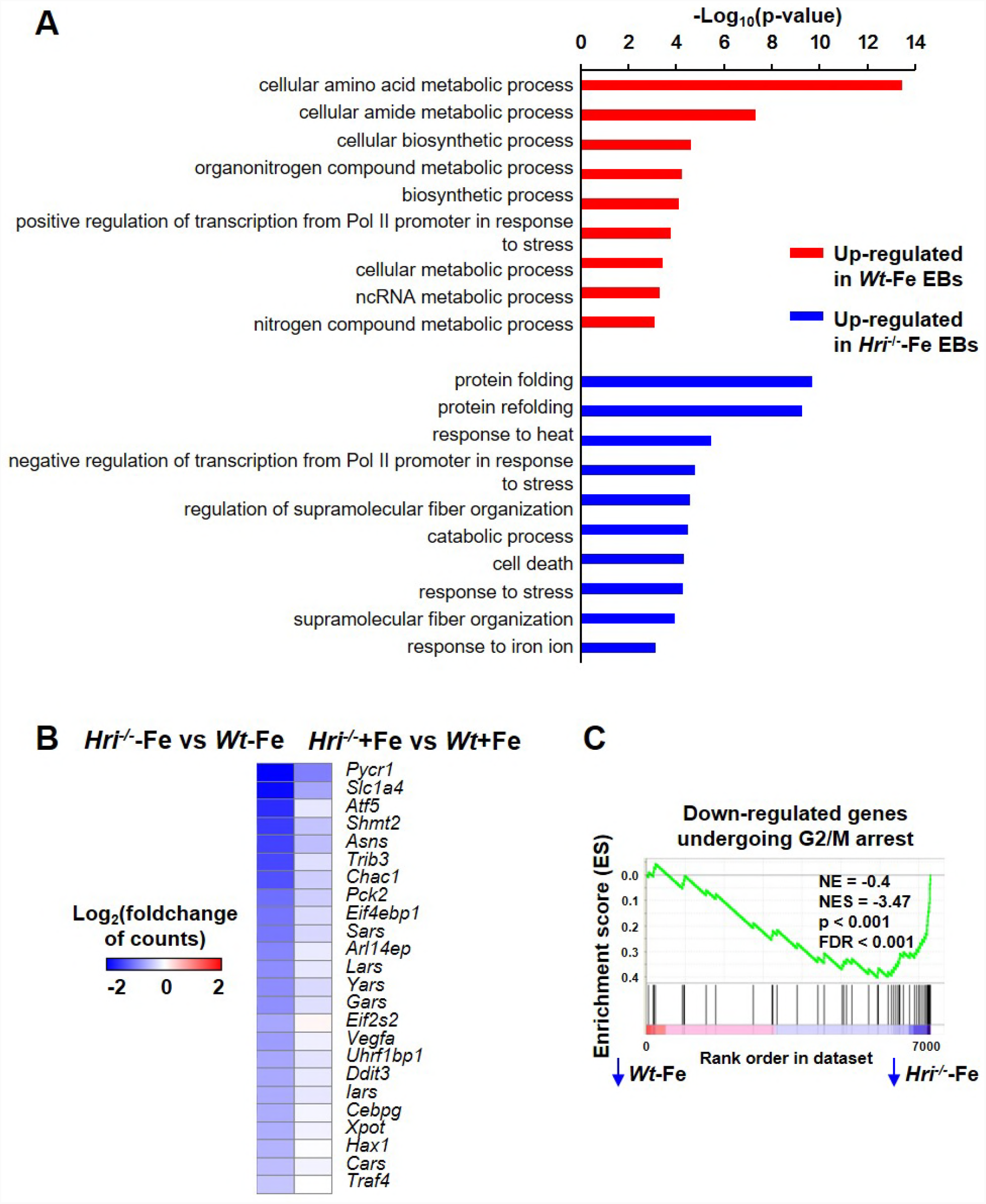
Analyses of differentially expressed mRNAs. GO analysis of significantly differentially expressed mRNAs between *Hri*^−/−^−Fe and *Wt*−Fe EBs. The heatmaps of differentially expressed ISR-target genes between *Hri*^−/−^−Fe and *Wt*−Fe EBs or between *Hri*^−/−^+Fe and *Wt*+Fe EBs. **(C)** Gene Set Enrichment Analysis revealed G2/M arrest pathway was downregulated in *Hri*^−/−^−Fe EBs as compared to *Wt*−Fe EBs.

**Figure supplement 5.**
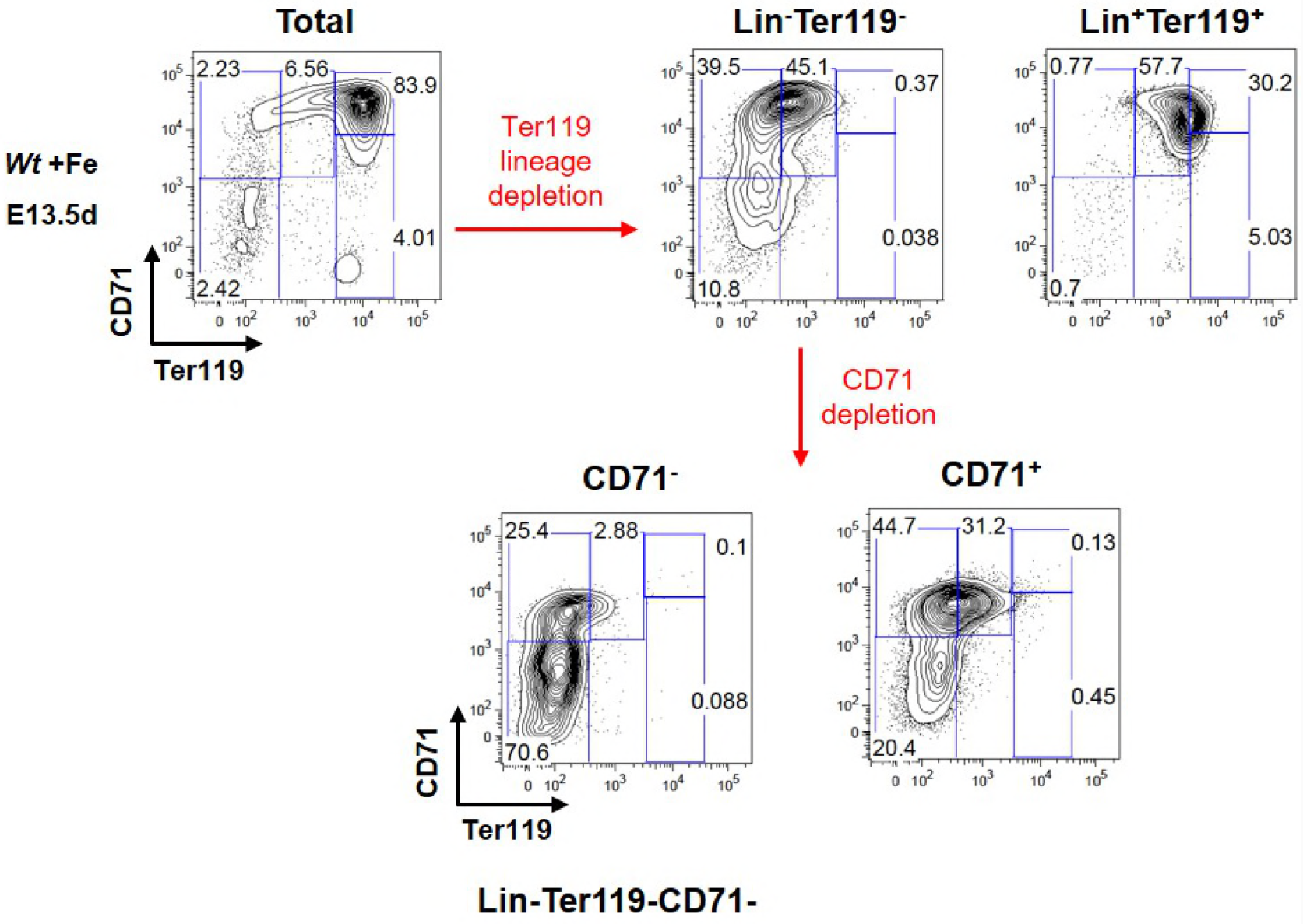
Enrichment of Lin^−^Ter119^−^CD71^−^ erythroid progenitors from *Wt*+Fe E13.5 FLs for *ex vivo* experiments.

**Figure supplement 6.**
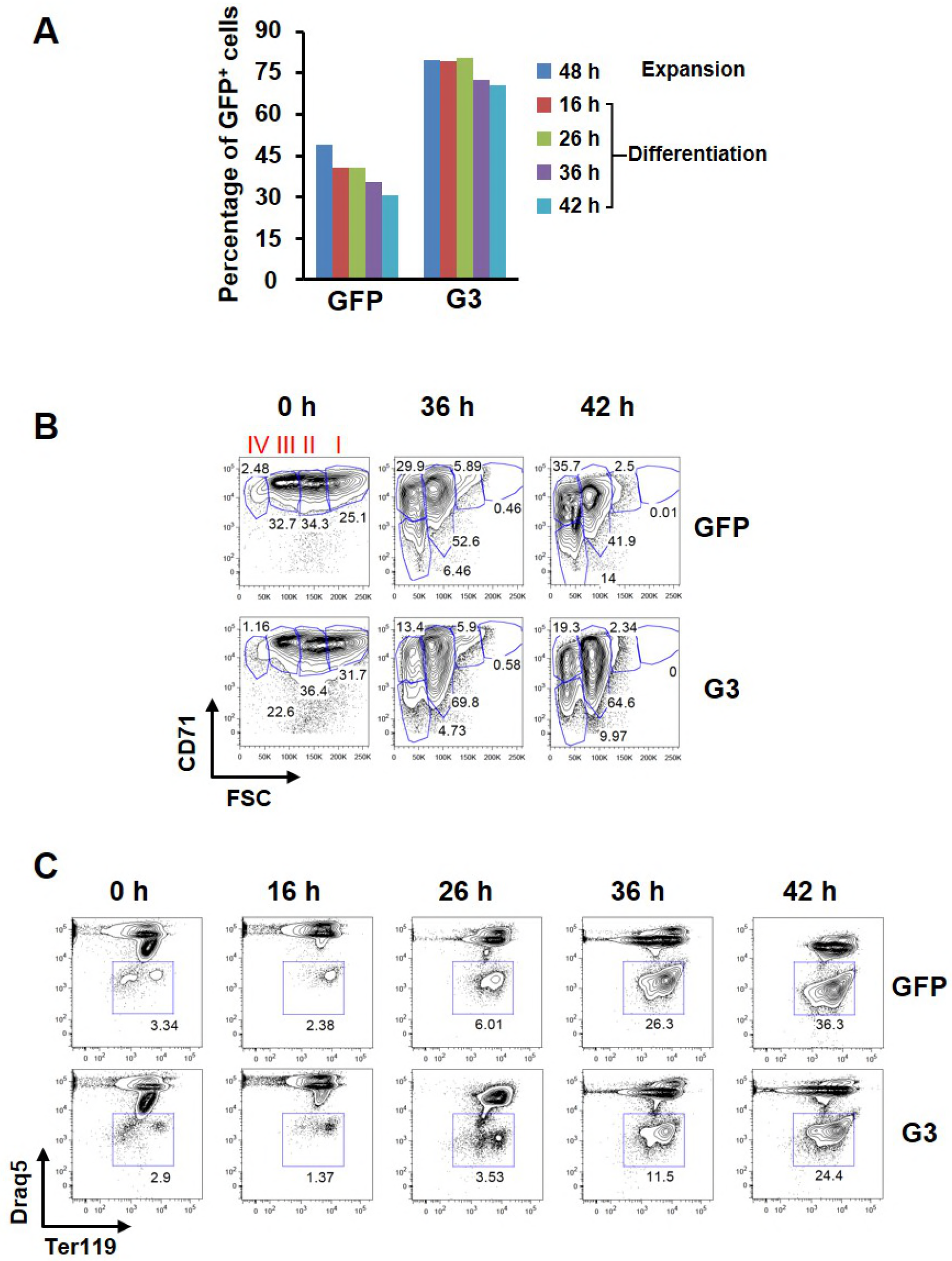
Effect of knockdown of *Grb10* on FL differentiation *ex vivo*. **(A)** Percentages of GFP^+^ cells during expansion and differentiation phases after infection of GFP control and *Grb10* shRNA_G3 recombinant retroviruses. **(B)** Representative plots of erythroid differentiation, corresponding to Figure 6D. **(C)** Enucleation of GFP^+^ cells during differentiation by flow cytometry using anti-Ter119 and Draq5 florescent dye.

**Figure supplement 7.**
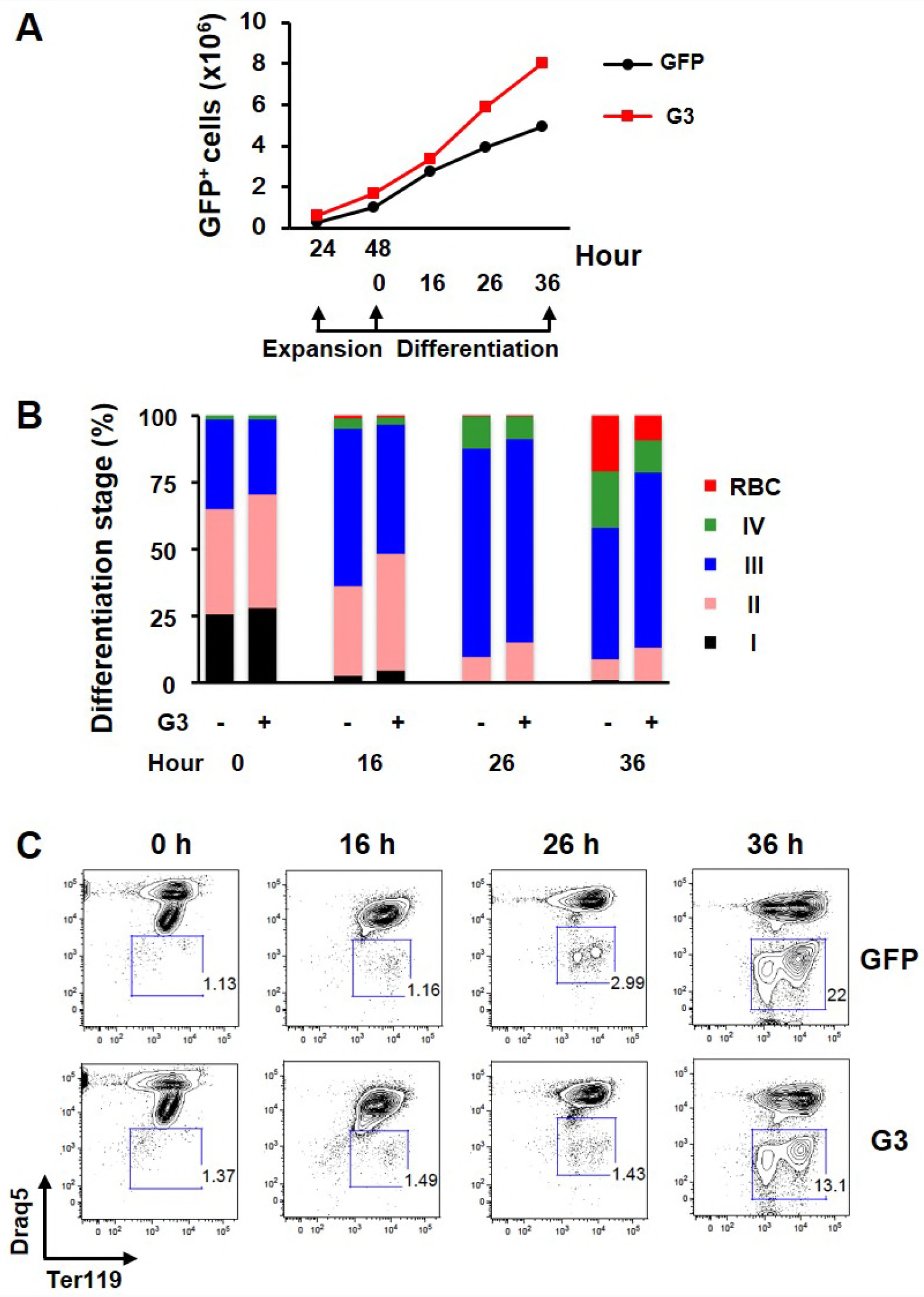
Proliferation and differentiation of FL erythroid progenitors upon *Grb10* knockdown. **(A)** The proliferation, **(B)** percentage of cells at erythroid differentiation stages, and **(C)** enucleation of GFP^+^ cells after infection of GFP control and *Grb10* shRNA_G3 recombinant retroviruses. These results were obtained from an independent separate experiment from the results shown in Figure 6.

## Table supplements

**Table Supplement 1.**
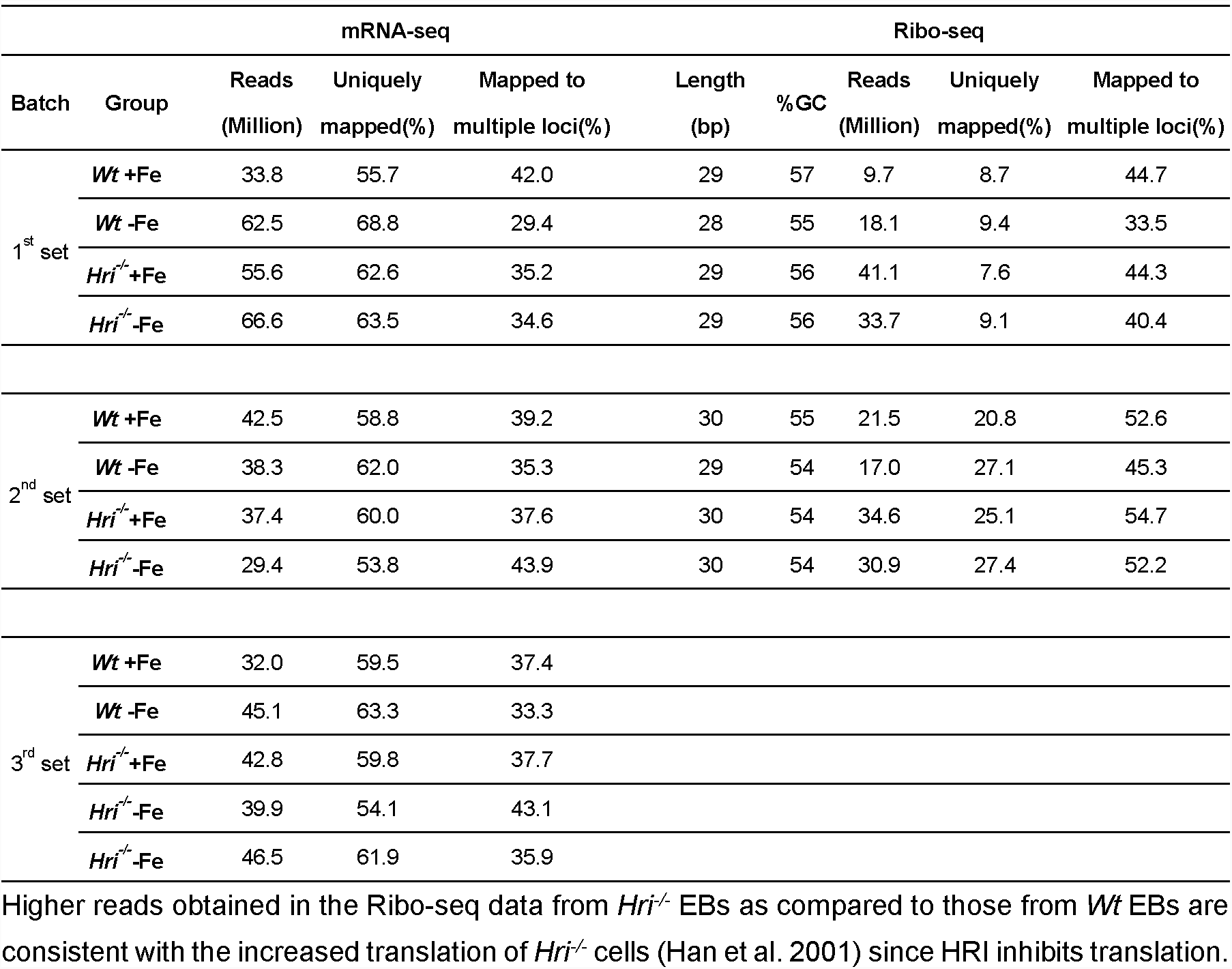
Quality control and mapping of Ribo-seq and mRNA-seq.

**Table Supplement 2 Complete gene list of differentially translated mRNAs**

As determined using xtail package of R/Bioconductor, genes with reads greater than 50 at least in one of the conditions in each comparison were included in TE (translational efficiency) calculation. NA indicates that gene was not included in TE calculation due to reads below cut-off.

**Table Supplement 3.** Higher ribosome occupancy at the AUG of uORF1 in *Atf4* mRNA.

| Gene Symbol | Type | Condition | Ribo-seq/mRNA-seq | *Atf4*/*eIF2s1* | *Atf4*/*Rps6* |
| --- | --- | --- | --- | --- | --- |
| *Atf4* | uORF1 | *Wt*+Fe | 10.7 | 5.7 | 21.3 |
|  |  | *Hri*<sup>-/-</sup>+Fe | 11.4 | 5.1 | 8.0 |
|  |  | *Wt*-Fe | 7.2 | 5.0 | 15.4 |
|  |  | *Hri*<sup>-/-</sup>-Fe | 20.3 | 12.8 | 13.5 |
| *eIF2s1* (*eIF2α*) | Canonical ORF | *Wt*+Fe | 1.9 |  |  |
|  |  | *Hri*<sup>-/-</sup>+Fe | 2.2 |  |  |
|  |  | *Wt*-Fe | 1.4 |  |  |
|  |  | *Hri*<sup>-/-</sup>-Fe | 1.6 |  |  |
| *Rps6* | Canonical ORF | *Wt*+Fe | 0.5 |  |  |
|  |  | *Hri*<sup>-/-</sup>+Fe | 1.4 |  |  |
|  |  | *Wt*-Fe | 0.5 |  |  |
|  |  | *Hri*<sup>-/-</sup>-Fe | 1.5 |  |  |

To calculate ribosome occupancy at AUG, Ribo-seq reads were normalized with the respective mRNA-seq reads. The AUG of uORF1 in *Atf4* mRNA was compared to the AUG of canonical ORFs of *eIF2s1* (*eIF2α*) and *Rps6*.

**Table Supplement 4.** No significant changes in TE of iron homeostasis-related genes.

| Gene Symbol | IRE | Log2(FoldChange of TE) |  |  |  |
| --- | --- | --- | --- | --- | --- |
|  |  | *Hri*<sup>-/-</sup>+Fe vs *Wt*+Fe | *Hri*<sup>-/-</sup>-Fe vs *Wt*-Fe | *Wt*-Fe vs *Wt*+Fe | *Hri*<sup>-/-</sup>-Fe vs *Hri*<sup>-/-</sup>+Fe |
| *Ftl1* | 5' UTR | 0.05 | 1.18 | -0.03 | 1.08 |
| *Fth1* | 5' UTR | -0.01 | 0.40 | -0.12 | 0.26 |
| *Alas2* | 5' UTR | -0.66 | 0.27 | -0.75 | 0.15 |
| *Aco2* | 5' UTR | -0.37 | -0.14 | 0.22 | 0.44 |
| *Slc40a1 (Fpn)* | 5' UTR | -0.47 | 0.56 | -0.39 | 0.61 |
| *Tfrc (Tfr1)* | 3' UTR | 0.01 | 0.29 | 0.07 | 0.32 |
| *Slc11a2 (Dmt1)* | 3' UTR | 0.15 | 0.14 | 0.21 | 0.18 |
| *Slc25a37 (Mfrn1)* | NA | 0.01 | 0.60 | -0.61 | -0.05 |
| *Ncoa4* | NA | -0.53 | 1.19 | -1.06 | 0.63 |
| *Tfr2* | NA | -0.97 | -0.16 | -0.48 | 0.31 |
| *Herc2* | NA | -1.89 | -1.08 | -0.92 | -0.14 |

NA indicates no iron-responsive element (IRE). UTR, untranslated region. TE, translational efficiency.

**Table supplement 5 Complete gene list of differentially expressed mRNAs**

As determined using DESeq2 package of R/Bioconductor, genes with the mean of reads of the conditions greater than 100 were used for further analysis.

Note: this table is a separate file in Excel format.

**Table Supplement 6.**
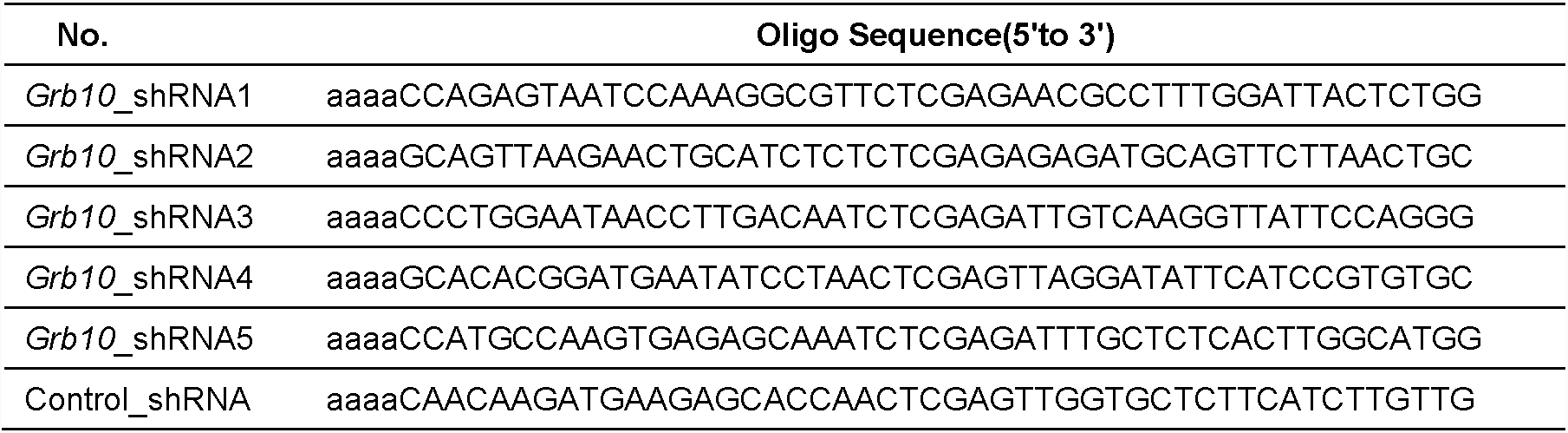
DNA sequences of shRNA oligonucleotides.

**Table Supplement 7.**
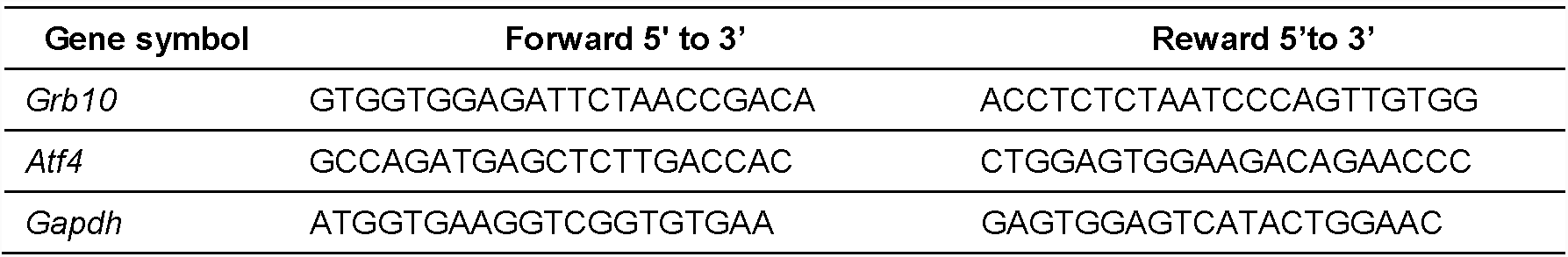
DNA sequence of qPCR primers.

**Table Supplement 8.** List of antibodies used for Western blot analysis.

| Symbol | Brand | Cat# |
| --- | --- | --- |
| ATF4 | Cell Signaling Technology | 11815 |
| GRB10 | Millipore | 07-2182 |
| HRI | JJ Chen Lab | Han *et al.*, 2001 |
| Cyclin D3 | Cell Signaling Technology | 2936 |
| pAKT (Ser473) | Cell Signaling Technology | 4060 |
| AKT | Cell Signaling Technology | 4691 |
| β-Actin | Cell Signaling Technology | 3700 |
| eIF2α | Cell Signaling Technology | 5324 |

## Method supplements

### Library preparations of Ribo-seq

To preserve polyribosomes, sorted EBs were washed twice with cold phosphate-buffered saline (PBS) and then treated with cycloheximde (100 μg/ml) at 37°C for 5 min, followed by isolation of RPFs as previously described (Guo et al. 2010, Ingolia, Lareau, and Weissman 2011) using ARTseq-Ribosome Profiling Kit (Illumina, San Diego, CA). Briefly, cells were lysed in cold polysome buffer on ice for 10 min. To generate RPFs, lysates were digested with Rnase 1. Monosomes were then purified by S-400 gel filtration spin columns (GE Healthcare, Chicago, IL). RNAs in monosomes were extracted, purified and followed by size selection of 28-34 nucleotides RPFs using denaturing polyacrylamide gel electrophoresis. Ribosomal RNA contaminants in the RPF preparations were removed using Ribo-Zero rRNA Removal Kit (Illumina, San Diego, CA). RPFs were then purified by 15% urea-polyacrylamide gel electrophoresis. Libraries of RPFs were prepared as described (Ingolia et al. 2009, Ingolia, Lareau, and Weissman 2011) for Illumina next generation DNA sequencing. RPFs were dephosphorylated and linkers were ligated using T4 RNA ligase. RNA samples were then reverse transcribed, circularized and PCR amplified for 12 cycles. PCR products were subjected to gel purification before sequencing. Results of Ribo-seq presented in Figure 2 were from the second set of experiments. For the first set of Ribo-seq experiment, EBs were treated with harringtonine (2 μg/ml) at 37°C for 2 min followed by cycloheximide treatment as described above. However, harringtonine treatment at this condition did not worked well for EBs as ribosome occupancy was observed throughout many mRNAs. For the second set of Ribo-seq experiment, harringtonine was not used. Nonetheless, increased translation of *Atf4, Pppr115a* and *Ddit3* mRNAs was also observed in both sets of Ribo-seq.

### Genome-wide data analysis

Raw data (fastq files) were trimmed by Cutadapt to remove adapters and reads with a base quality score less than 10. Reads with length less than 26 or 9 nucleotides were also discarded for Ribo-seq and mRNA-seq samples, respectively. For Ribo-seq samples, reads containing rRNA and tRNA sequences were further removed using Bowtie2. After quality control analysis using FastQC, reads were mapped to mouse genome mm10 (UCSC) using STAR aligner with maxima of 2 mismatches and 8 multiple loci. Quality of Ribo-seq data was examined by the triplet periodicity using RibORF (Ji et al. 2015).

For the visualization of ribosome occupancies, all mapped reads (bam files) that overlap each bin (bin size = 1) were firstly calculated and normalized to effective mouse genome size to get a 1x depth of coverage (RPGC) using bamCoverage of deepTools (Ramirez et al. 2016). Then, ribosome occupancies were visualized using the Integrative Genomics Viewer (Broad Institute of Harvard and MIT). Since no significant change at mRNA levels among four conditions, only mapped reads of *Wt*+Fe EBs were shown for mRNA-seq data in Figure 2C-D, Figure 3C, Figure supplement 1B-C and Figure supplement 2A.

The uniquely mapped reads were counted using HTseq for gene coverage analysis, transcript per million (TPM) calculation, translational efficiency (TE) calculation and analysis of differentially expressed mRNAs (DEG). Genes with uniquely mapped reads greater than 25 in at least one of the conditions were used for gene coverage analysis and TPM calculation.

Generally, RPFs are piled up around the start codons and stop codons due to the slower kinetics of translation initiation and termination. The use of cycloheximide to freeze polysomes during the preparation of RPFs also enhances the pile up of reads near start codons. We, therefore, removed the first 15 and the last 5 codons from reads counting for the TE calculation of Ribo-seq data, in which TE is defined as the ratio of RPF counts to mRNA counts. TE was calculated using xtail package of R/Bioconductor, with parameter ‘bins = 10000’, to determine differentially translated genes between different conditions (Xiao et al. 2016). Genes with reads greater than 50 in at least one of the conditions were included in each comparison and were used for TE calculation (Table supplement 2). Genes with adjusted p-value < 0.05 together with log_2_(foldchange of TE) > 1.5 or < −1.5 were considered as significantly differentially translated in Figure 2A while adjusted p-value < 0.05 together with log_2_(foldchange of TE) > 1 or < −1 were considered as significantly differentially translated in Figure 2B, Figure 3A, Figure 3D and Figure supplement 1A.

For Gene Set Enrichment Analysis (GSEA), pre-ranked gene lists were obtained using formula −Log_10_(adjusted p-value) × Log_2_(foldchange of TE). Mouse gene symbols were converted into human symbols using biomaRt package of R/Bioconductor. GSEA was conducted using GSEA tool from Broad Institute with c2.all.v6.0.symbols.gmt database, which includes 4738 gene sets. Gene sets shown in Figure 3B were REACTOME_FORMATION_OF_THE_TERNARY_COMPLEX_AND_SUBSEQUENTLY _THE_43S_COMPL (rank 13), KEGG_RIBOSOME (rank 1) and BILANGES_SERUM_AND_RAPAMYCIN_SENSITIVE_GENES (rank 9). Gene set shown in Figure supplement 4C was BHATI_G2M_ARREST_BY_2METHOXYESTRADIOL_DN (rank 17).

Differentially expressed genes from mRNA-seq data were determined using DESeq2 package of R/Bioconductor, and genes with the mean of reads of all conditions greater than 100 were used for further analysis (Table supplement 5). Genes with adjusted p-value < 0.05 were considered as significantly differentially expressed in Figure 4 and Figure supplement 4A-B. Lists of 5’ TOP/TOP-like genes were obtained from Thoreen *et al* (Thoreen et al. 2012) while the integrated stress response (ISR) gene list was derived from Palam *et al* (Palam et al. 2015).

Gene ontology analysis was performed using g:Profiler (Reimand et al. 2016) for TE and DEG data sets. The START App was employed to generate the volcano plots, in which red dots indicate significantly differentially translated or expressed genes, while green and gray dots indicate non significantly changed genes (Nelson et al. 2017). Heatmaps were plotted using GENE-E tool from the Broad Institute.

### Purification of FL erythroid progenitors for *ex vivo* culture and differentiation

In brief, total FL cells were mechanically dissociated by pipetting in PBS containing 2% FBS (Atlanta Biologicals, Norcross, GA), 2.5 mM EDTA and 10 mM glucose. Cells were labeled with biotin-conjugated anti-B220, -CD3, -CD11b, -CD11c, -GR1, -Ter119 (BioLegend, San Diego, CA) and -CD41 (eBioscience, San Diego, CA) antibodies. Lineage negative (Lin^−^Ter119^−^) cells were subjected to a second purification step to obtain Lin^−^Ter119^−^CD71^−^ erythroid progenitors using biotin-conjugated anti-CD71 antibodies (BioLegend, San Diego, CA and BD Biosciences, San Jose, CA).

Purified FL erythroid progenitors were expanded in StemPro-34 medium complemented with 10% supplement, 2mM L-glutamine, 1% Penicillin-Streptomycin (P-S), 10^−4^ M β-mercaptoethanol, 10^−6^ M dexamethasone, 0.5 U/mL Epo, 100 ng/mL mouse Stem Cell Factor (mSCF), and then differentiated in Iscove modified Dulbecco medium (IMDM) containing 10% FCS (fetal calf serum, Gemini Bio-Products, West Sacramento, CA), 10% plasma derived serum (Animal Technologies, Tyler, TX), 2 mM L-glutamine, 1% P-S, 10^−4^ M β-mercaptoethanol, 5 U/mL Epo.

## References

An, X., V. P. Schulz, J. Li, K. Wu, J. Liu, F. Xue, J. Hu, N. Mohandas, and P. G. Gallagher. 2014. “Global transcriptome analyses of human and murine terminal erythroid differentiation.” Blood 123 (22):3466–77. doi:10.1182/blood-2014-01-548305.

Anderson, Sheila A, Christopher P Nizzi, Yuan-I. Chang, Kathryn M Deck, Paul J Schmidt, Bruno Galy, Alisa Damnernsawad, Aimee T Broman, Christina Kendziorski, Matthias W Hentze, Mark D Fleming, Jing Zhang, and Richard S Eisenstein. 2013. “The IRP1-HIF-2α Axis Coordinates Iron and Oxygen Sensing with Erythropoiesis and Iron Absorption.” Cell Metabolism 17 (2):282–290. doi:https://doi.org/10.1016/j.cmet.2013.01.007.

Bellelli, R., G. Federico, A. Matte, D. Colecchia, A. Iolascon, M. Chiariello, M. Santoro, L. De Franceschi, and F. Carlomagno. 2016. “NCOA4 Deficiency Impairs Systemic Iron Homeostasis.” Cell Rep 14 (3):411–421. doi:10.1016/j.celrep.2015.12.065.

Brush, M. H., D. C. Weiser, and S. Shenolikar. 2003. “Growth arrest and DNA damage-inducible protein GADD34 targets protein phosphatase 1 alpha to the endoplasmic reticulum and promotes dephosphorylation of the alpha subunit of eukaryotic translation initiation factor 2.” Mol Cell Biol 23 (4):1292–303.

Charalambous, Marika, Florentia M. Smith, William R. Bennett, Tracey E. Crew, Francesca Mackenzie, and Andrew Ward. 2003. “Disruption of the imprinted Grb10 gene leads to disproportionate overgrowth by an Igf2-independent mechanism.” Proceedings of the National Academy of Sciences 100 (14):8292.

Chen, J. J. 2007. “Regulation of protein synthesis by the heme-regulated eIF2alpha kinase: relevance to anemias.” Blood 109 (7):2693–9. doi:10.1182/blood-2006-08-041830.

Chen, J. J. 2014. “Translational control by heme-regulated eIF2alpha kinase during erythropoiesis.” Curr Opin Hematol 21 (3):172–8. doi:10.1097/MOH.0000000000000030.

Chiabrando, D., S. Mercurio, and E. Tolosano. 2014. “Heme and erythropoieis: more than a structural role.” Haematologica 99 (6):973–83. doi:10.3324/haematol.2013.091991.

Connor, J. H., D. C. Weiser, S. Li, J. M. Hallenbeck, and S. Shenolikar. 2001. “Growth arrest and DNA damage-inducible protein GADD34 assembles a novel signaling complex containing protein phosphatase 1 and inhibitor 1.” Mol Cell Biol 21 (20):6841–50. doi:10.1128/mcb.21.20.6841-6850.2001.

Cooperman, Sharon S., Esther G. Meyron-Holtz, Hayden Olivierre-Wilson, Manik C. Ghosh, Joseph P. McConnell, and Tracey A. Rouault. 2005. “Microcytic anemia, erythropoietic protoporphyria, and neurodegeneration in mice with targeted deletion of iron-regulatory protein 2.” Blood 106 (3):1084.

Galy, Bruno, Dunja Ferring, Belen Minana, Oliver Bell, Heinz G. Janser, Martina Muckenthaler, Klaus Schümann, and Matthias W. Hentze. 2005. “Altered body iron distribution and microcytosis in mice deficient in iron regulatory protein 2 (IRP2).” Blood 106 (7):2580.

Ghosh, Manik C., Wing-Hang Tong, Deliang Zhang, Hayden Ollivierre-Wilson, Anamika Singh, Murali C. Krishna, James B. Mitchell, and Tracey A. Rouault. 2008. “Tempol-mediated activation of latent iron regulatory protein activity prevents symptoms of neurodegenerative disease in IRP2 knockout mice.” Proceedings of the National Academy of Sciences 105 (33):12028.

Ghosh, Manik C, De-Liang Zhang, Suh Young Jeong, Gennadiy Kovtunovych, Hayden Ollivierre-Wilson, Audrey Noguchi, Tiffany Tu, Thomas Senecal, Gabrielle Robinson, Daniel R Crooks, Wing-Hang Tong, Kavitha Ramaswamy, Anamika Singh, Brian B Graham, Rubin M Tuder, Zu-Xi Yu, Michael Eckhaus, Jaekwon Lee, Danielle A Springer, and Tracey A Rouault. 2013. “Deletion of Iron Regulatory Protein 1 Causes Polycythemia and Pulmonary Hypertension in Mice through Translational Derepression of HIF2α.” Cell Metabolism 17 (2):271–281. doi:https://doi.org/10.1016/j.cmet.2012.12.016.

Grevet, J. D., X. Lan, N. Hamagami, C. R. Edwards, L. Sankaranarayanan, X. Ji, S. K. Bhardwaj, C. J. Face, D. F. Posocco, O. Abdulmalik, C. A. Keller, B. Giardine, S. Sidoli, B. A. Garcia, S. T. Chou, S. A. Liebhaber, R. C. Hardison, J. Shi, and G. A. Blobel. 2018. “Domain-focused CRISPR screen identifies HRI as a fetal hemoglobin regulator in human erythroid cells.” Science 361 (6399):285–290. doi:10.1126/science.aao0932.

Guo, Huili, Nicholas T. Ingolia, Jonathan S. Weissman, and David P. Bartel. 2010. “Mammalian microRNAs predominantly act to decrease target mRNA levels.” Nature 466:835. doi:10.1038/nature09267 https://www.nature.com/articles/nature09267#supplementary-information.

Hahn, C. K., and C. H. Lowrey. 2013. “Eukaryotic initiation factor 2alpha phosphorylation mediates fetal hemoglobin induction through a post-transcriptional mechanism.” Blood 122 (4):477–85. doi:10.1182/blood-2013-03-491043.

Han, A. P., M. D. Fleming, and J. J. Chen. 2005. “Heme-regulated eIF2alpha kinase modifies the phenotypic severity of murine models of erythropoietic protoporphyria and beta-thalassemia.” J Clin Invest 115 (6):1562–70. doi:10.1172/JCI24141.

Han, A. P., C. Yu, L. Lu, Y. Fujiwara, C. Browne, G. Chin, M. Fleming, P. Leboulch, S. H. Orkin, and J. J. Chen. 2001. “Heme-regulated eIF2alpha kinase (HRI) is required for translational regulation and survival of erythroid precursors in iron deficiency.” Embo j 20 (23):6909–18. doi:10.1093/emboj/20.23.6909.

Harding, H. P., Y. Zhang, H. Zeng, I. Novoa, P. D. Lu, M. Calfon, N. Sadri, C. Yun, B. Popko, R. Paules, D. F. Stojdl, J. C. Bell, T. Hettmann, J. M. Leiden, and D. Ron. 2003. “An integrated stress response regulates amino acid metabolism and resistance to oxidative stress.” Mol Cell 11 (3):619–33.

Hinnebusch, Alan G. 1996. “Translational Control of Gcn4: Gene-Specific Regulation by Phosphorylation of Eif2.” In Translational Control, edited by J.W.B. Hershey, M.B. Mathews and N. Sonenberg, 199-244. Cold Spring Harbor: Cold Spring Harbor Laboratory Press.

Hu, W., B. Yuan, and H. F. Lodish. 2014. “Cpeb4-mediated translational regulatory circuitry controls terminal erythroid differentiation.” Dev Cell 30 (6):660–72. doi:10.1016/j.devcel.2014.07.008.

Igarashi, K., and J. Sun. 2006. “The heme-Bach1 pathway in the regulation of oxidative stress response and erythroid differentiation.” Antioxid Redox Signal 8 (1-2):107–18. doi:10.1089/ars.2006.8.107.

Ingolia, N. T., G. A. Brar, N. Stern-Ginossar, M. S. Harris, G. J. Talhouarne, S. E. Jackson, M. R. Wills, and J. S. Weissman. 2014. “Ribosome profiling reveals pervasive translation outside of annotated protein-coding genes.” Cell Rep 8 (5):1365–79. doi:10.1016/j.celrep.2014.07.045.

Ingolia, N. T., S. Ghaemmaghami, J. R. Newman, and J. S. Weissman. 2009. “Genome-wide analysis in vivo of translation with nucleotide resolution using ribosome profiling.” Science 324 (5924):218–23. doi:10.1126/science.1168978.

Ingolia, N. T., L. F. Lareau, and J. S. Weissman. 2011. “Ribosome profiling of mouse embryonic stem cells reveals the complexity and dynamics of mammalian proteomes.” Cell 147 (4):789–802. doi:10.1016/j.cell.2011.10.002.

Ji, Zhe, Ruisheng Song, Aviv Regev, and Kevin Struhl. 2015. “Many lncRNAs, 5’UTRs, and pseudogenes are translated and some are likely to express functional proteins.” eLife 4:e08890. doi:10.7554/eLife.08890.

Kaufman, Randal J. 2000. “Double-Stranded Rna-Activated Protein Kinase Pkr.” In Translational Control of Gene Expression, edited by J.W.B. Hershey, N. Sonenberg and M.B. Matthews, 503–28. Cold Spring Harbor: Cold Spring Harbor Laboratory Press.

Kerenyi, M. A., and S. H. Orkin. 2010. “Networking erythropoiesis.” J Exp Med 207 (12):2537–41. doi:10.1084/jem.20102260.

Khajuria, R. K., M. Munschauer, J. C. Ulirsch, C. Fiorini, L. S. Ludwig, S. K. McFarland, N. J. Abdulhay, H. Specht, H. Keshishian, D. R. Mani, M. Jovanovic, S. R. Ellis, C. P. Fulco, J. M. Engreitz, S. Schutz, J. Lian, K. W. Gripp, O. K. Weinberg, G. S. Pinkus, L. Gehrke, A. Regev, E. S. Lander, H. T. Gazda, W. Y. Lee, V. G. Panse, S. A. Carr, and V. G. Sankaran. 2018. “Ribosome Levels Selectively Regulate Translation and Lineage Commitment in Human Hematopoiesis.” Cell. doi:10.1016/j.cell.2018.02.036.

Kingsley, P. D., E. Greenfest-Allen, J. M. Frame, T. P. Bushnell, J. Malik, K. E. McGrath, C. J. Stoeckert, and J. Palis. 2013. “Ontogeny of erythroid gene expression.” Blood 121 (6):e5–e13. doi:10.1182/blood-2012-04-422394.

Kojima, E., A. Takeuchi, M. Haneda, A. Yagi, T. Hasegawa, K. Yamaki, K. Takeda, S. Akira, K. Shimokata, and K. Isobe. 2003. “The function of GADD34 is a recovery from a shutoff of protein synthesis induced by ER stress: elucidation by GADD34-deficient mice.” Faseb j 17 (11):1573–5. doi:10.1096/fj.02-1184fje.

Liu, S., S. Bhattacharya, A. Han, R. N. Suragani, W. Zhao, R. C. Fry, and J. J. Chen. 2008. “Haem-regulated eIF2alpha kinase is necessary for adaptive gene expression in erythroid precursors under the stress of iron deficiency.” Br J Haematol 143 (1):129–37. doi:10.1111/j.1365-2141.2008.07293.x.

Lopez, Anthony, Patrice Cacoub, Iain C. Macdougall, and Laurent Peyrin-Biroulet. 2016. “Iron deficiency anaemia.” The Lancet 387 (10021):907–916. doi:10.1016/s0140-6736(15)60865-0.

Mancias, J. D., L. Pontano Vaites, S. Nissim, D. E. Biancur, A. J. Kim, X. Wang, Y. Liu, W. Goessling, A. C. Kimmelman, and J. W. Harper. 2015. “Ferritinophagy via NCOA4 is required for erythropoiesis and is regulated by iron dependent HERC2-mediated proteolysis.” Elife 4. doi:10.7554/eLife.10308.

Mancias, J. D., X. Wang, S. P. Gygi, J. W. Harper, and A. C. Kimmelman. 2014. “Quantitative proteomics identifies NCOA4 as the cargo receptor mediating ferritinophagy.” Nature 509 (7498):105–9. doi:10.1038/nature13148.

Masuoka, H. C., and T. M. Townes. 2002. “Targeted disruption of the activating transcription factor 4 gene results in severe fetal anemia in mice.” Blood 99 (3):736–45.

Mills, E. W., J. Wangen, R. Green, and N. T. Ingolia. 2016. “Dynamic Regulation of a Ribosome Rescue Pathway in Erythroid Cells and Platelets.” Cell Rep 17 (1):1–10. doi:10.1016/j.celrep.2016.08.088.

Muckenthaler, Martina U., Stefano Rivella, Matthias W. Hentze, and Bruno Galy. 2017. “A Red Carpet for Iron Metabolism.” Cell 168 (3):344–361. doi:10.1016/j.cell.2016.12.034.

Nelson, Jonathan W., Jiri Sklenar, Anthony P. Barnes, and Jessica Minnier. 2017. “The START App: a web-based RNAseq analysis and visualization resource.” Bioinformatics 33 (3):447–449. doi:10.1093/bioinformatics/btw624.

Novoa, I., H. Zeng, H. P. Harding, and D. Ron. 2001. “Feedback inhibition of the unfolded protein response by GADD34-mediated dephosphorylation of eIF2alpha.” J Cell Biol 153 (5):1011–22.

Pakos-Zebrucka, Karolina, Izabela Koryga, Katarzyna Mnich, Mila Ljujic, Afshin Samali, and Adrienne M. Gorman. 2016. “The integrated stress response.” EMBO reports 17 (10):1374–1395. doi:10.15252/embr.201642195.

Palam, L. R., J. Gore, K. E. Craven, J. L. Wilson, and M. Korc. 2015. “Integrated stress response is critical for gemcitabine resistance in pancreatic ductal adenocarcinoma.” Cell Death & Disease 6:e1913. doi:10.1038/cddis.2015.264 https://www.nature.com/articles/cddis2015264#supplementary-information.

Paolini, Nahuel A., Kat S. Moore, Franca M. di Summa, Ivo F. A. C. Fokkema, Peter A. C. ‘t Hoen, and Marieke von Lindern. 2018. “Ribosome profiling uncovers selective mRNA translation associated with eIF2 phosphorylation in erythroid progenitors.” PLOS ONE 13 (4):e0193790. doi:10.1371/journal.pone.0193790.

Patterson, A. D., M. C. Hollander, G. F. Miller, and A. J. Fornace, Jr. 2006. “Gadd34 requirement for normal hemoglobin synthesis.” Mol Cell Biol 26 (5):1644–53. doi:10.1128/mcb.26.5.1644-1653.2006.

Pavitt, G. D., and D. Ron. 2012. “New insights into translational regulation in the endoplasmic reticulum unfolded protein response.” Cold Spring Harb Perspect Biol 4 (6). doi:10.1101/cshperspect.a012278.

Plasschaert, R. N., and M. S. Bartolomei. 2015. “Tissue-specific regulation and function of Grb10 during growth and neuronal commitment.” Proc Natl Acad Sci U S A 112 (22):6841–7. doi:10.1073/pnas.1411254111.

Ramirez, F., D. P. Ryan, B. Gruning, V. Bhardwaj, F. Kilpert, A. S. Richter, S. Heyne, F. Dundar, and T. Manke. 2016. “deepTools2: a next generation web server for deep-sequencing data analysis.” Nucleic Acids Res 44 (W1):W160–5. doi:10.1093/nar/gkw257.

Reimand, J., T. Arak, P. Adler, L. Kolberg, S. Reisberg, H. Peterson, and J. Vilo. 2016. “g:Profiler-a web server for functional interpretation of gene lists (2016 update).” Nucleic Acids Res 44 (W1):W83–9. doi:10.1093/nar/gkw199.

Ron, David, and Heather P. Harding. 2000. “Perk and Translational Control by Stress in Endoplasmic Reticulum.” In Translational Control of Gene Expression, edited by J.W.B. Hershey, N. Sonenberg and M.B. Matthews, 547–60. Cold Springs Harbor: Cold Spring Harbor Laboratory Press.

Sankaran, V. G., L. S. Ludwig, E. Sicinska, J. Xu, D. E. Bauer, J. C. Eng, H. C. R. A. Metcalf, Y. Natkunam, S. H. Orkin, P. Sicinski, E. S. Lander, and H. F. Lodish. 2012. “Cyclin D3 coordinates the cell cycle during differentiation to regulate erythrocyte size and number.” Genes Dev 26 (18):2075–87. doi:10.1101/gad.197020.112.

Sankaran, Vijay G., and Stuart H. Orkin. 2013. “The Switch from Fetal to Adult Hemoglobin.” Cold Spring Harbor Perspectives in Medicine 3 (1):a011643. doi:10.1101/cshperspect.a011643.

Scheuner, D., B. Song, E. McEwen, C. Liu, R. Laybutt, P. Gillespie, T. Saunders, S. Bonner-Weir, and R. J. Kaufman. 2001. “Translational control is required for the unfolded protein response and in vivo glucose homeostasis.” Mol Cell 7 (6):1165–76.

Schranzhofer, M., M. Schifrer, J. A. Cabrera, S. Kopp, P. Chiba, H. Beug, and E. W. Mullner. 2006. “Remodeling the regulation of iron metabolism during erythroid differentiation to ensure efficient heme biosynthesis.” Blood 107 (10):4159–67. doi:10.1182/blood-2005-05-1809.

Singh, Seema, Arvind Dev, Rakesh Verma, Anamika Pradeep, Pradeep Sathyanarayana, Jennifer M. Green, Aishwarya Narayanan, and Don M. Wojchowski. 2012. “Defining an EPOR-Regulated Transcriptome for Primary Progenitors, including Tnfr-sf13c as a Novel Mediator of EPO-Dependent Erythroblast Formation.” PLOS ONE 7 (7):e38530. doi:10.1371/journal.pone.0038530.

Smith, F. M., L. J. Holt, A. S. Garfield, M. Charalambous, F. Koumanov, M. Perry, R. Bazzani, S. A. Sheardown, B. D. Hegarty, R. J. Lyons, G. J. Cooney, R. J. Daly, and A. Ward. 2007. “Mice with a disruption of the imprinted Grb10 gene exhibit altered body composition, glucose homeostasis, and insulin signaling during postnatal life.” Mol Cell Biol 27 (16):5871–86. doi:10.1128/mcb.02087-06.

Stonestrom, A. J., S. C. Hsu, K. S. Jahn, P. Huang, C. A. Keller, B. M. Giardine, S. Kadauke, A. E. Campbell, P. Evans, R. C. Hardison, and G. A. Blobel. 2015. “Functions of BET proteins in erythroid gene expression.” Blood 125 (18):2825–34. doi:10.1182/blood-2014-10-607309.

Suragani, R. N., R. S. Zachariah, J. G. Velazquez, S. Liu, C. W. Sun, T. M. Townes, and J. J. Chen. 2012. “Heme-regulated eIF2alpha kinase activated Atf4 signaling pathway in oxidative stress and erythropoiesis.” Blood 119 (22):5276–84. doi:10.1182/blood-2011-10-388132.

Thom, C. S., E. A. Traxler, E. Khandros, J. M. Nickas, O. Y. Zhou, J. E. Lazarus, A. P. Silva, D. Prabhu, Y. Yao, C. Aribeana, S. Y. Fuchs, J. P. Mackay, E. L. Holzbaur, and M. J. Weiss. 2014. “Trim58 degrades Dynein and regulates terminal erythropoiesis.” Dev Cell 30 (6):688–700. doi:10.1016/j.devcel.2014.07.021.

Thoreen, C. C., L. Chantranupong, H. R. Keys, T. Wang, N. S. Gray, and D. M. Sabatini. 2012. “A unifying model for mTORC1-mediated regulation of mRNA translation.” Nature 485 (7396):109–13. doi:10.1038/nature11083.

Ulirsch, J. C., J. N. Lacy, X. An, N. Mohandas, T. S. Mikkelsen, and V. G. Sankaran. 2014. “Altered chromatin occupancy of master regulators underlies evolutionary divergence in the transcriptional landscape of erythroid differentiation.” PLoS Genet 10 (12):e1004890. doi:10.1371/journal.pgen.1004890.

Wang, L., B. Balas, C. Y. Christ-Roberts, R. Y. Kim, F. J. Ramos, C. K. Kikani, C. Li, C. Deng, S. Reyna, N. Musi, L. Q. Dong, R. A. DeFronzo, and F. Liu. 2007. “Peripheral disruption of the Grb10 gene enhances insulin signaling and sensitivity in vivo.” Mol Cell Biol 27 (18):6497–505. doi:10.1128/mcb.00679-07.

Wilkinson, Nicole, and Kostas Pantopoulos. 2013. “IRP1 regulates erythropoiesis and systemic iron homeostasis by controlling HIF2α mRNA translation.” Blood 122 (9):1658.

Xiao, Zhengtao, Qin Zou, Yu Liu, and Xuerui Yang. 2016. “Genome-wide assessment of differential translations with ribosome profiling data.” Nature Communications 7:11194. doi:10.1038/ncomms11194 https://www.nature.com/articles/ncomms11194#supplementary-information.

Yan, X., H. A. Himburg, K. Pohl, M. Quarmyne, E. Tran, Y. Zhang, T. Fang, J. Kan, N. J. Chao, L. Zhao, P. L. Doan, and J. P. Chute. 2016. “Deletion of the Imprinted Gene Grb10 Promotes Hematopoietic Stem Cell Self-Renewal and Regeneration.” Cell Rep 17 (6):1584–1594. doi:10.1016/j.celrep.2016.10.025.

Zhang, Shuping, Alejandra Macias-Garcia, Jason Velazquez, Elena Paltrinieri, Randal J. Kaufman, and Jane-Jane Chen. 2018. “HRI coordinates translation by eIF2aP and mTORC1 to mitigate ineffective erythropoiesis in mice during iron deficiency.” Blood 131 (4):450.

